# Blood cells of adult *Drosophila* do not expand, but control survival after bacterial infection by induction of *Drosocin* around their reservoir at the respiratory epithelia

**DOI:** 10.1101/578864

**Authors:** Pablo Sanchez Bosch, Kalpana Makhijani, Leire Herboso, Katrina S Gold, Rowan Baginsky, Katie J Woodcock, Brandy Alexander, Katelyn Kukar, Sean Corcoran, Debra Ouyang, Corinna Wong, Elodie JV Ramond, Christa Rhiner, Eduardo Moreno, Bruno Lemaitre, Frederic Geissmann, Katja Brückner

**Author notes:** Corresponding Author and Lead Contact: 35 Medical Center Way San Francisco, CA 94143-0669. Equal contribution, in reverse alphabetical order (Pablo Sanchez Bosch, Kalpana Makhijani, Leire Herboso, Katrina S Gold, Rowan Baginsky).

## Abstract

*Drosophila melanogaster* has been an excellent model for innate immunity, but the role and regulation of adult blood cells and organismal immunity have remained incompletely understood. Here we address these questions in a comprehensive investigation of the blood cell system in adult *Drosophila*. As a central finding, we reveal the largest reservoir of blood cells (hemocytes) at the respiratory epithelia (tracheal air sacs) and fat body of the thorax and head. We show that most hemocytes of adult *Drosophila* are phagocytic macrophages (plasmatocytes), derived by more than 60% from the embryonic lineage that parallels vertebrate tissue macrophages. Surprisingly, in contrast to hemocytes at the larval stage, we find no capacity of the adult blood cell system to expand. Instead, we demonstrate its central role in relaying an innate immune response to tissues surrounding the blood cell reservoir: Hemocytes, through Imd signaling and the Jak/Stat pathway ligand Upd3, act as sentinels of bacterial infection that induce expression of the antimicrobial peptide gene *Drosoci*n in the respiratory epithelia and colocalizing domains of the fat body. We demonstrate that endogenous *Drosocin* expression in these tissues promotes animal survival after bacterial infection. Our work identifies the first molecular step in a new relay of organismal immunity, establishing adult *Drosophila* as model to dissect mechanisms of inter-organ immunity.

## Introduction

*Drosophila melanogaster* has greatly promoted our understanding of innate immunity and blood cell development, but the capacity of the adult animal as a model remains a matter of debate. Most studies reported a lack of new blood cell production (Lanot et al., 2001; Mackenzie et al., 2011; Woodcock et al., 2015) and immunosenescence (Felix et al., 2012; Mackenzie et al., 2011), while a recent publication claimed continued hematopoietic activity in adult *Drosophila* (Ghosh et al., 2015).

*Drosophila* blood cells, or hemocytes, emerge from two lineages that persist into adult life, showing parallels with the two myeloid systems in vertebrates (Gold and Brückner, 2014, 2015; Holz et al., 2003). First, hemocytes originating in the embryo parallel vertebrate tissue macrophages, as they quickly differentiate into plasmatocytes (macrophage-like cells), and subsequently proliferate extensively, mainly in the hematopoietic pockets (HPs) of the larva (Gold and Brückner, 2014, 2015; Makhijani et al., 2011; Makhijani and Brückner, 2012). At least some embryonic-lineage plasmatocytes can further differentiate into other blood cell types such as crystal cells and, under immune challenge, lamellocytes (Bretscher et al., 2015; Gold and Brückner, 2015; Leitao and Sucena, 2015; Makhijani et al., 2011; Markus et al., 2009). Second, hemocytes originating in the lymph gland (LG) also give rise to plasmatocytes, crystal cells and lamellocytes, yet in the lymph gland they are predominantly generated via differentiation from blood cell progenitors (prohemocytes) (Banerjee et al., 2019; Gold and Brückner, 2015; Jung et al., 2005; Letourneau et al., 2016). At the beginning of metamorphosis, hemocytes from both the hematopoietic pockets and the lymph gland enter the open circulatory system, leading to intermixing of the two blood cell lineages. (Gold and Brückner, 2015; Grigorian et al., 2011; Lanot et al., 2001; Makhijani et al., 2011). The subsequent fate and capacity of the adult blood cells is an ongoing matter of debate. To clarify this question, we dedicated the first part of our study to comprehensively investigate the hematopoietic capacity of the blood cell system in adult *Drosophila*.

For the second part of the study, we focused on the role of the adult blood cell pool in the humoral immune response, identifying a new system of organismal innate immunity in *Drosophila*. Historically, *Drosophila* has been instrumental in the discovery of innate immunity and Toll like receptor (TLR) signaling (Lemaitre and Hoffmann, 2007). Toll- and the related Immune Deficiency (Imd) signaling, are evolutionary conserved NFкB family pathways that have been studied in detail regarding their upstream activation by pathogens and other inputs, and downstream signal transduction components and mechanisms (Lemaitre and Hoffmann, 2007). Targets include antimicrobial peptides (AMPs), which have been investigated for their transcriptional gene regulation and functional properties (Lemaitre and Hoffmann, 2007; Zasloff, 2002). TLR signaling has been well established also in vertebrate systems for its roles in infection and inflammation (Beutler, 2009; Kopp and Medzhitov, 1999; Takeda and Akira, 2005). However, despite the detailed knowledge of TLR signaling and innate immunity at the cellular and molecular level, it has been far less understood how multiple tissues or organs communicate with each other to elicit local innate immune responses.

Addressing these two major questions, our study comprehensively illuminates basic principles of the blood cell system in adult *Drosophila* and it role in multi-tissue organismal immunity. We identify an extensive, previously unknown blood cell reservoir at the respiratory epithelia and fat body, investigate its dynamics, and probe for various signs of hematopoiesis. We demonstrate a key role of adult blood cells as sentinels of bacterial infection that trigger a humoral response in their reservoir, i.e. the respiratory epithelia and colocalizing domains of the fat body. This response culminates in the expression of the AMP gene *Drosocin*, which we show is significant for animal survival after bacterial infection. We identifiedy a requirement for Imd signaling and Upd3 expression in hemocytes as a first molecular step in this relay of organismal immunity, laying the foundation for the use of adult *Drosophila* to dissect additional mechanisms of multi-tissue innate immunity in the future.

## Results

### The respiratory epithelia are the largest reservoir of blood cells in adult Drosophila

To investigate the blood cell system in adult *Drosophila*, we started out by examining the anatomical sites of hemocytes. We visualized blood cells by fluorescent labeling with macrophage (plasmatocyte)-specific *Hemolectin HmlΔ-GAL4* (Sinenko and Mathey-Prevot, 2004) driving *UAS-GFP*, or direct reporters *HmlΔ-DsRed* or *HmlΔDsRednls* (Makhijani et al., 2011). To gain an unbiased overview of hemocyte locations throughout the animal, we took a cryosectioning approach. In addition, we imaged hemocytes through the cuticle of whole flies. Hemocytes are largely resident (sessile), and show consistent enrichment in specific areas. Surprisingly, we found that the largest pool of hemocytes colonizes the respiratory epithelia, in particular the extensive air sacs of the thorax and head (Fig. 1A-E). This location was identified in our cryosections by comparison with anatomical features of the respiratory epithelia (Manning and Krasnow, 1993; Whitten, 1957), and the unique blue autofluorescence of the respiratory epithelia when exposed to UV light (Kim et al., 2012). Localization of hemocytes with the respiratory epithelia was further confirmed by colabeling with the tracheal driver *btl-GAL4* expressing *UAS-GFP* (Guha and Kornberg, 2005) (Fig. 1D, Suppl.Fig. 1C). In intact flies, hemocyte localization at the respiratory epithelia is visible around the eyes and posterior head, and in the thorax laterally, and dorsally near the wing hinges and scutellum (Suppl. Fig. 1A,C,D).

**Figure 1.**
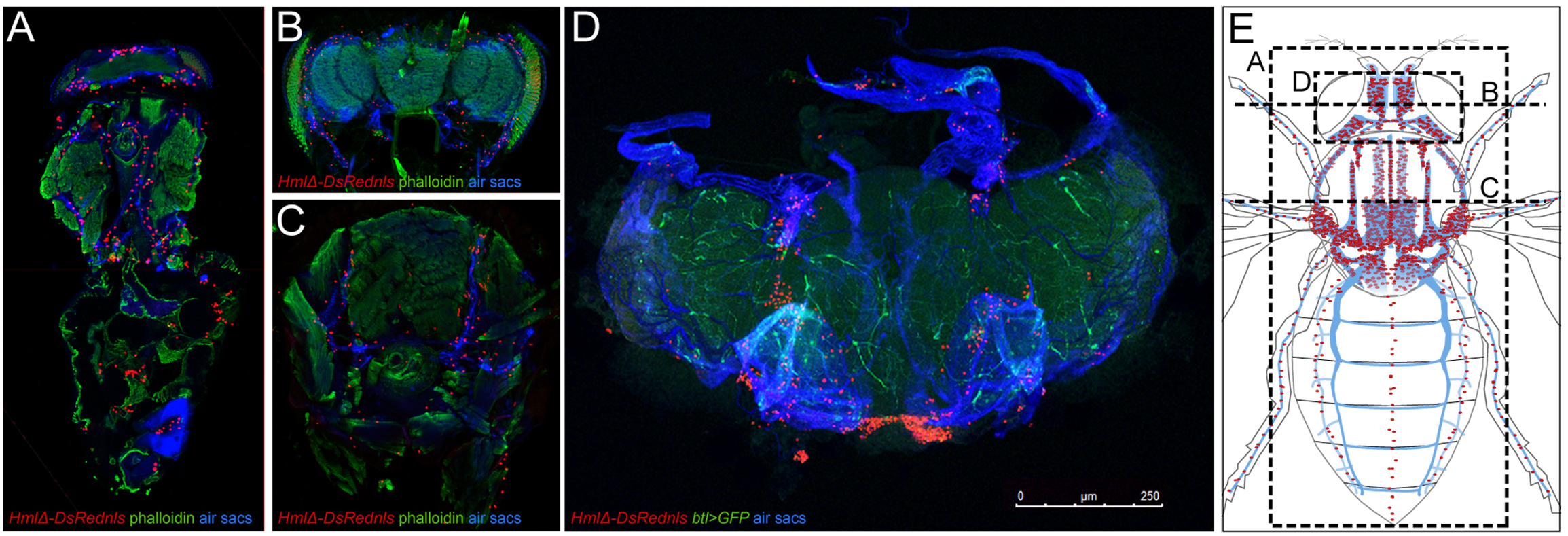
The respiratory epithelia provide the largest reservoir of hemocytes in adult *Drosophila*. (A-C) Cryosections of adult *Drosophila, HmlΔ-DsRednls* hemocytes red, phalloidin green, respiratory epithelia (air sacs) blue. (A) Longitudinal section, anterior up; (B) cross section of head, dorsal up; (C) cross section of thorax, dorsal up. (D) Adult *Drosophila*, genotype *HmlΔ-DsRed*/+; *btl-GAL4, UAS-GFP/+*; hemocytes red, tracheal marker green; respiratory epithelia (air sacs) blue; dissection of head, anterior up; size bar 250µm. (E) Schematics of respiratory epithelia (tracheal air sacs) of the thorax and head and other parts of the tracheal system in blue), hemocytes in red. Dashed lines indicate sections and dissected area shown in A-D. Note that model omits heart area, which is not visible in longitudinal section in (A). For full model including heart area, see Suppl. Fig. 1.

Consistent with previous reports (Dionne et al., 2003; Elrod-Erickson et al., 2000; Ghosh et al., 2015), we also saw a smaller fraction of hemocytes surrounding the heart (Suppl.Fig. 1A,B,D), accumulating in clusters at the ostia (Suppl. Fig.1B), which are the intake valves of the heart toward the open circulatory system. Quantification of hemocytes released from flies split into two parts, at the boundary of the thorax and abdomen, confirmed that the majority of hemocytes in adult *Drosophila* is located in the head and thorax (see Fig. 3F). Overall we conclude that, in adult *Drosophila*, the respiratory epithelia of the head and thorax provide the major reservoir of blood cells, which is distinct from smaller clusters of hemocytes at the ostia of the heart.

**Figure 2.**
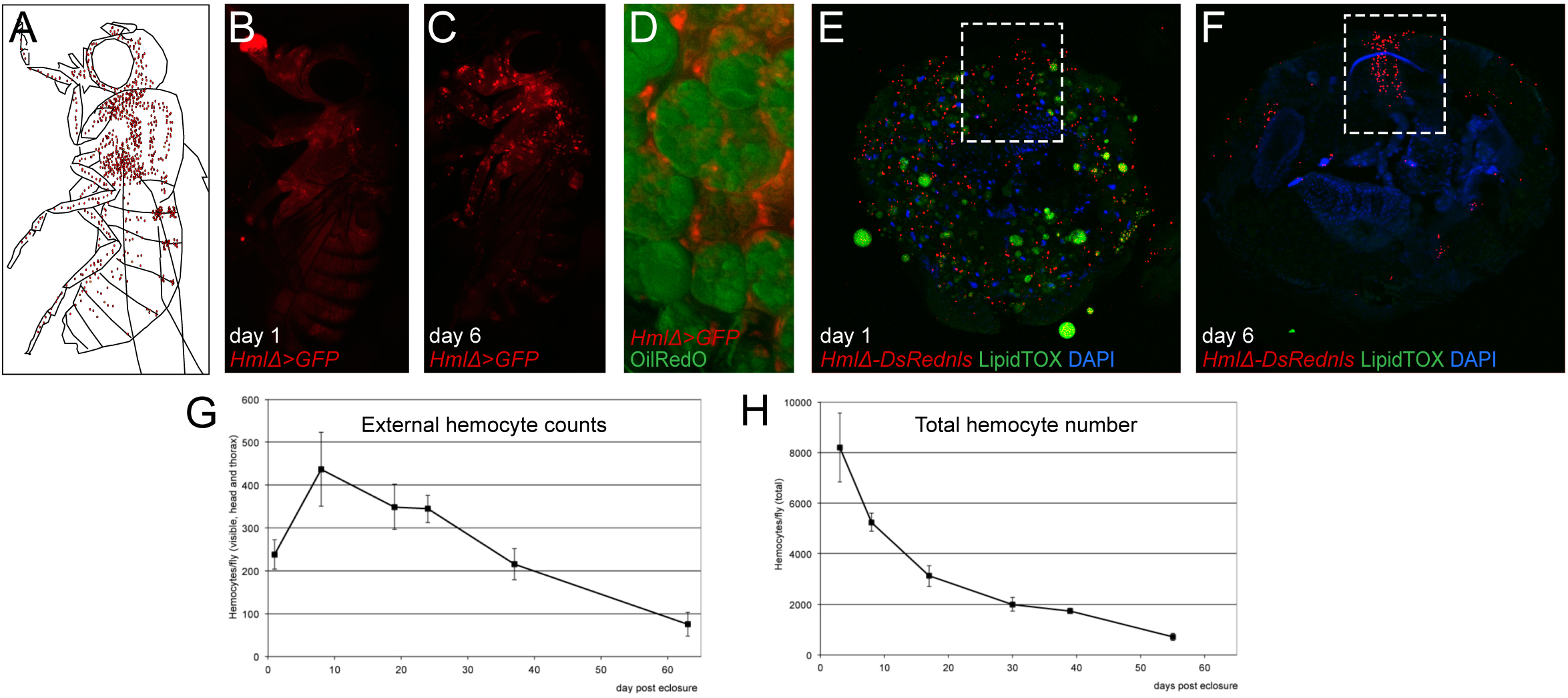
Developmental changes of hemocytes. (A-C) Lateral view of adult *Drosophila*, (A) Model, hemocytes red, (B,C) *HmlΔ-GAL4, UAS-GFP* (hemocytes, red pseudo color), (B) day 1 post eclosure, (C) day 6 post eclosure. (D) Larval fat body cells (Oil Red O pseudo-colored in green) with associated hemocytes (*HmlΔ-GAL4, UAS-GFP* pseudo-colored in red), dissection of abdomen. (E, F) cross sections of anterior abdomen, *HmlΔ-DsRednls* (hemocytes red), LipidTox (green), DAPI (blue); dashed box marks heart region; (E) day 1 post eclosure; (F) day 6 post eclosure. (G) External hemocyte quantification, time course; fluorescently labeled hemocytes that can be visually recognized through the cuticle were counted (see Methods). (H) Total hemocyte counts per animal, time course; all hemocytes were released by scraping and counted ex vivo.

**Figure 3.**
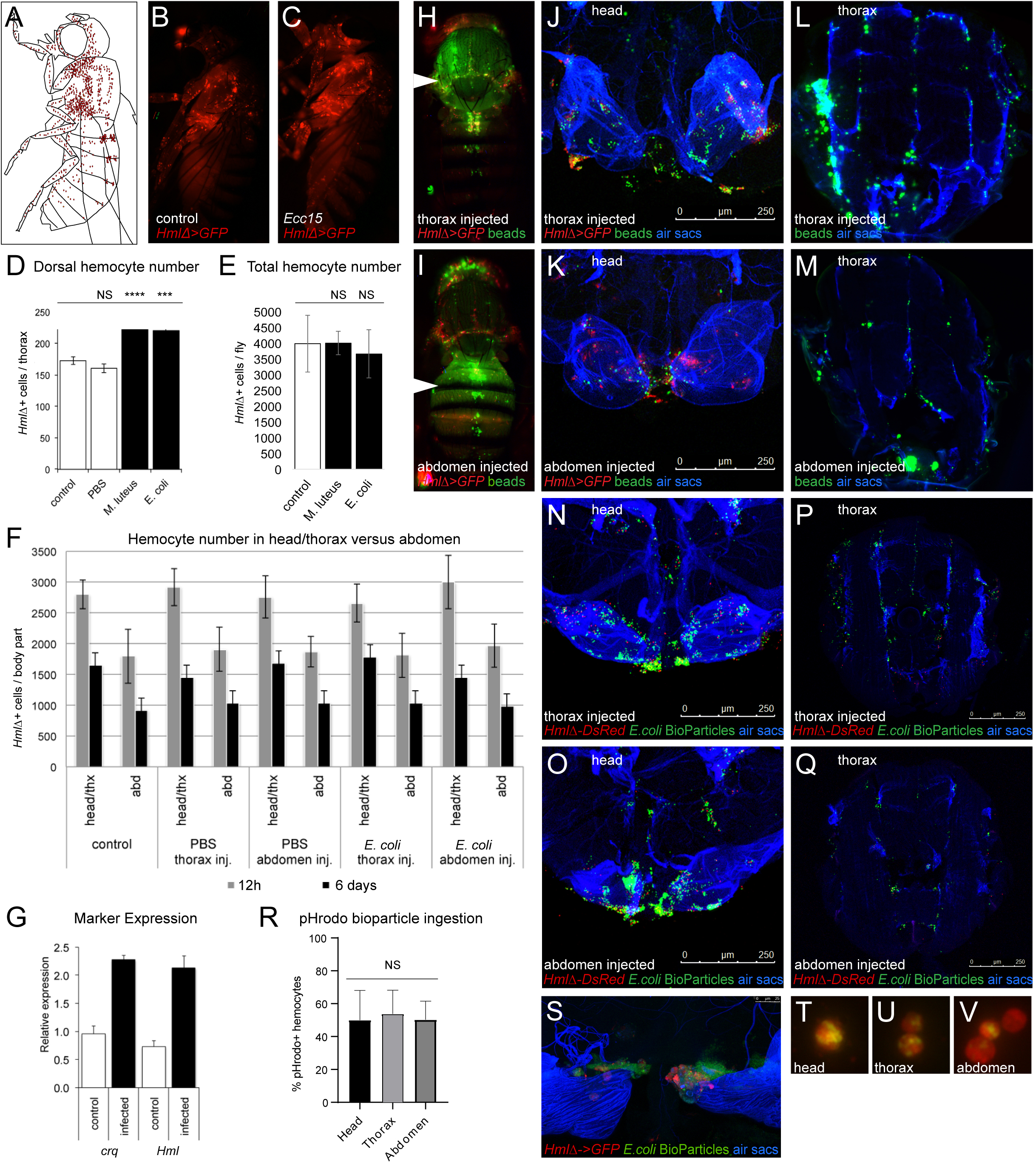
Infection-induced changes of hemocytes and accumulation of particles at the respiratory epithelia. (A-C) Lateral view of adult *Drosophila*; (A) Model, hemocytes red, (B, C) *HmlΔ-GAL4, UAS-GFP* (hemocytes red pseudo color), (B) no infection control, (C) *Ecc15* injection. (D) External hemocyte quantification, hemocyte number of the dorsal thorax and anterior abdomen, controls and injected flies as indicated; p values of paired 2-tailed t test are shown, *,**,***,or **** corresponding to p≤0.05, 0.01, 0.001, or 0.0001. (E) Total hemocyte counts, control and injected flies as indicated, all hemocytes were released by scraping. Average and standard deviation; 2-tailed t test shows no statistically significant difference (NS). (F) Total hemocyte counts of flies split in two parts, head and thorax versus abdomen; control and injected flies as indicated. Flies were injected into thorax or abdomen at 5 days post eclosion and assayed at 12h and 6d post injection. Average and standard deviation are shown. (G) qPCR expression levels of *Hml* and *Crq* from whole flies, −/+ infection, 48h post infection. 8 day old adults were injected with *E.coli* at OD 2. (H-M) Injection of fluorescent microbeads (green pseudocolor), *HmlΔ-GAL4, UAS-GFP* (hemocytes, red pseudocolor), respiratory epithelia (air sacs, blue). (H, J, L) Injection in thorax, (H) external view, (J) head dissection, (L) thorax cross section. (I, K, M) Injection in abdomen, (I) external view, (K) head dissection, (M) thorax cross section. (N-Q) Injection of fluorescent *E. coli* bioparticles (green), *HmlΔ-DsRed* for head dissections or *HmlΔ-DsRednls* for cryosections (hemocytes, red), respiratory epithelia (air sacs, blue). (N, P) Injection in thorax, (N) head dissection, (P) thorax cross section. (O, Q) Injection in abdomen, (O) head dissection, (Q) thorax cross section. (R) Injection of pHrodo *E. coli* bioparticles into *HmlΔ-DsRed* adults; percentages of hemocytes that phagocytosed pHrodo bioparticles from dissected head, thorax and abdomen 4 hours after injection. Mean with standard deviation and one-way ANOVA significance show no statistically significant difference (NS). (S) Dissected respiratory epithelia of the head from adults *HmlΔ-DsRed* (hemocytes, red) injected with pHrodo *E. coli* bioparticles (green) 4 hours after injection. (T-V) Examples of hemocytes with incorporated pHrodo bioparticles isolated from head, thorax, abdomen, corresponding to (R).

### Hemocytes relocate during maturation of adult Drosophila

Next we investigated the developmental timing of hemocyte localization to the respiratory system and heart. Newly eclosed *Drosophila* expressing a fluorescent plasmatocyte reporter show a diffuse glow in live imaging, which over the following 5-7 days develops into a more defined hemocyte pattern (Fig. 2B-C); in these mature adults, hemocytes then remain rather stationary over time (Suppl. Fig2A-C and (Woodcock et al., 2015)). While the visual change of hemocytes in young adults may suggest an increase in total hemocyte numbers, we actually found a different process to be underlying this phenomenon. Specifically, we discovered a major redistribution of hemocytes within the first days of adult life. Dissection and lipid dye staining of newly eclosed adults illustrates that hemocytes are attached to dissociated larval fat body cells (Fig. 2D) (Nelliot et al., 2006), forming a large mass throughout the abdomen and other parts of the fly (Fig. 2E). Around 5-7 days into adult life, cytolysis of the larval fat body cells is completed, allowing hemocytes to relocate to resident sites at the respiratory system and heart (Fig. 2F). Therefore, counting local hemocytes, e.g. in the heart area, gives the false impression of an increase in blood cells during the first days of adult life (Fig. 2E-F dashed box). An impression of increasing blood cell numbers is also given when counting fluorescently labeled hemocytes that can be visually recognized through the external cuticle (Fig. 2G). In contrast, when assessing hemocytes in cryosections (Fig. 2E-F), or quantifying total hemocytes from dissected adult flies, we discovered a continuous decline in hemocytes over various time points, even during the first week of adult life (Fig. 2H and (Woodcock et al., 2015)).

In summary, we find no evidence for a significant increase in total hemocyte numbers during adult maturation. Instead, we observe a dramatic redistribution of existing hemocytes during the first week of adult life; once larval fat body cells have cytolysed, hemocytes are free to move toward the periphery to their final destinations, with the majority of hemocytes colonizing the respiratory epithelia.

### Hemocytes do not expand after septic injury

Next we investigated whether bacterial infection could have an effect on hemocytes of the respiratory system and the heart (Fig. 3B, C). Injection of adult flies with gram-negative *Escherichia coli (E.coli), Enterobacter cloacae (E. cloacae), Erwinia carotovora carotovora 15* (*Ecc15*), or gram-positive *Micrococcus luteus (M. luteus)* resulted in increased hemocyte numbers in external counts 6 days post infection (Fig. 3D). Surprisingly, we discovered that absolute hemocyte numbers, which were quantified by blood cell release ex vivo, did not increase upon infection (Fig. 3E). Asking whether hemocytes may redistribute upon infection, we assessed ex vivo blood cell counts from flies split into 2 parts consisting of head plus thorax, versus abdomen (Fig. 3F). Similar as in maturation, bacterial immune challenge did not affect absolute hemocyte numbers in the two sections; likewise, there was no effect by the site of injection in the thorax or abdomen (Fig. 3F). Tracking down what could be the basis for the seemingly increased hemocyte appearance after infection, we found a rather unexpected explanation. Using qPCR analysis, we saw that bacterial infection leads to transcriptional upregulation of macrophage-specific *Hml* and other commonly used marker genes such as *croquemort (crq)* (Fig. 3G), a phenomenon that actually has been described before (De Gregorio et al., 2001; Franc et al., 1999). This suggests that increased *HmlΔ* reporter fluorescence after bacterial infection likely accounts for elevated GFP expression, increasing the number of hemocytes that can be visually recognized through the cuticle, while absolute hemocyte counts are not raised.

To further address the role of the respiratory hemocyte reservoirs in infection, we examined the dynamics of particle accumulation, injecting fluorescently labeled microbeads or *E. coli* bioparticles (Molecular Probes/Invitrogen). Independent of the site of injection in the thorax or abdomen, microbeads or bioparticles quickly accumulate along the respiratory epithelia of the thorax and head (Fig. 3H-Q), and at the ostia of the heart (Fig. 3H,I), matching typical sites of hemocyte residence. Comparable localization of bioparticles was observed under hemocyte ablation conditions (Suppl. Fig. 3B), suggesting that this accumulation pattern is independent of hemocytes or their phagocytic activity. Hemocytes at the respiratory epithelia and other locations are capable of phagocytosis, as we confirmed by ingestion of injected bioparticles marked with regular label (Fig. 3S) or pHrodo, a pH dependent dye highly fluorescent in acidic compartments such as lysosomes following phagocytic ingestion (Suppl. Fig. 3C). Fractions of pHrodo positive hemocytes were similar when released from head, thorax or abdomen of split flies (Fig. 3R, T-V), suggesting comparable phagocytic activity and similar ratios of hemocyte- and particle accumulation in all body parts.

In summary, we find major accumulation of foreign particles in the hemocyte reservoirs lining the respiratory epithelia of *Drosophila*. This localization is independent of the presence of phagocytes, suggesting passive transport through the hemolymph and physical retention in these areas. Bacterial infection does not substantially change the overall localization of hemocytes, and does not cause significant increase in the total number of hemocytes per animal. Instead, we find a substantial increase in the expression of macrophage markers, which likely accounts for the impression of increased hemocyte numbers as they appear though the cuticle.

### The majority of adult hemocytes derive from the embryonic lineage

Next we sought to determine the ontogenesis of the adult blood cell system. Blood cells in adult *Drosophila* are known to derive from two lineages (Holz et al., 2003), the embryonic lineage that parallels tissue macrophages, and the lymph gland lineages that parallels progenitor-based blood cell production (Fig. 4A) (Banerjee et al., 2019; Gold and Brückner, 2014, 2015). To address their relative contributions to the adult animal, we used a flipout-*lacZ* lineage tracing approach (Makhijani et al., 2011; Weigmann and Cohen, 1999). In this approach, spatially and temporally controlled Flp recombinase induces reconstitution and permanent expression of a lacZ reporter transgene, which can be followed and compared relative to other cell markers. First, we used the method to mark early embryonic hemocytes (under control of *srpHemo-Gal4*) and examined animals at the pupal and adult stage (Fig. 4B-E). Indeed, this approach showed many *lacZ* positive hemocytes in the pupa (Fig. 4D,D’) and in the adult, including hemocytes attached to larval fat body cells in newly eclosed flies (Fig. 4E). We obtained similar results when we labeled differentiated embryonic-lineage plasmatocytes in the young larva using *HmlΔ-GAL4* (Fig. 4F-I). The method also allowed to quantify the contribution of embryonic-lineage hemocytes to the adult blood cell pool. Comparing the fraction of *lacZ* positive cells among plasmatocytes in the adult, relative to those in the 3^rd^ instar larva, we estimate that more than 60% of adult macrophages originate from the embryonic lineage (Fig. 4I). This is surprising, given the previously prevailing view that the majority of adult hemocytes would derive from the lymph gland (Lanot et al., 2001). Based on our findings we expect that less than 40% of adult hemocytes derive from the lymph gland lineage, although we can only infer this contribution as the relatively weak expression of early lymph gland GAL4 drivers left our lineage tracing attempts of the lymph gland unsuccessful.

**Figure 4.**
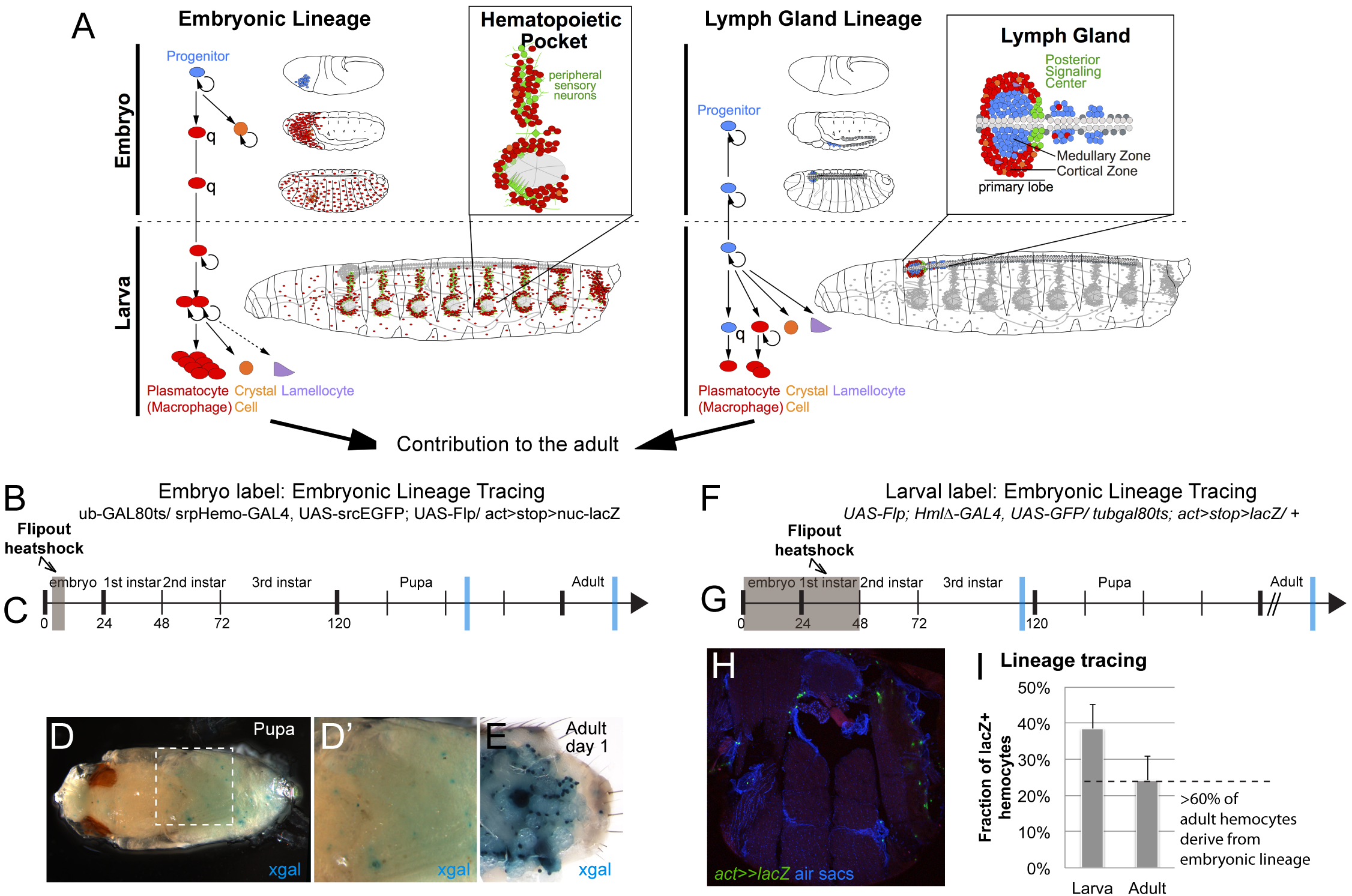
Contribution of the two hemocyte lineages to the adult blood cell pool. (A) Timeline of the two blood cell lineages in *Drosophila*, i.e. the embryonic and lymph gland lineage, with the major sites of hematopoiesis in the larva (hematopoietic pockets and lymph gland). Both lineages persist into the adult but the relative contribution has remained unclear. (B-E) *flipout-lacZ* lineage tracing using embryonically expressed *srpHemo-GAL4*. (B) Experimental genotype *tub-GAL80*^*ts*^ */ srpHemo-GAL4, UAS-srcEGFP; UAS-Flp/ act>stop>nuc-lacZ*. (C) Timeline of induction, hemocytes of the embryo were labeled in a 6h time window of Flp expression (grey box); blue bars mark time points of samples shown in (D-E). (D-D’) *lacZ*/βGal positive hemocytes in the pupa, x-gal staining (blue); (E) *lacZ*/βGal positive hemocytes in the dissected abdomen of an adult, x-gal staining (blue); note that adult staining appears stronger due to decreased permeability of whole mount pupae, individual variation of lineage tracing, and occasional labeling of larval fat body cells. (F-I) *flipout-lacZ* lineage tracing of embryonic-lineage hemocytes in the larva, using *HmlΔ-GAL4*. (F) Experimental genotype *UAS-Flp; HmlΔ-GAL4, UAS-GFP/ tub-GAL80*^*ts*^*; act>stop>lacZ/ +.* (G) timeline, induction at 0-48h AEL (grey box). Note that at this stage lymph gland hemocytes do not express *HmlΔ-GAL4* and therefore remain unlabeled; *HmlΔ-GAL4* is expressed in embryonic-lineage hemocytes from late embryonic stage onward. Blue bars mark time points of samples examined and quantified (see H, I); (H) Thorax cross section of adult fly, genotype as in (F), *lacZ*/βGal positive hemocytes green (anti-βGal), air sacs and DAPI in blue. (I) Fraction of *lacZ*/βGal positive hemocytes relative to all Crq positive hemocytes in late 3^rd^ instar larvae and in the adult; relative to each other, these numbers suggest contribution of the embryonic lineage to more that 60% of the adult blood cell pool (dashed line).

Taken together, we conclude that the majority of adult blood cells derive from the embryonic hemocyte lineage that proliferates in the larva and resembles tissue macrophages in vertebrates (Gold and Brückner, 2014, 2015; Makhijani et al., 2011; Makhijani and Brückner, 2012).

### Adult Drosophila show no signs of new hemocyte production

Given that adult hemocytes originate from embryonic-lineage hemocytes, which retain the ability to proliferate as macrophages in the larva (Makhijani et al., 2011), and the lymph gland hemocyte lineage, which may comprise undifferentiated progenitors with putative proliferation potential that arise in the posterior lobes (Grigorian et al., 2011; Jung et al., 2005), we wanted to investigate the proliferative capacity and differentiation status of adult hemocytes. First we used in vivo EdU (5-ethynyl-2’-deoxyuridine) incorporation into newly synthesized DNA to detect proliferation of macrophages and their putative progenitors. Labeling all EdU-incorporating cells over a continuous time period of two weeks and examining the entire pool of adult hemocytes ex vivo, we surprisingly did not find any EdU positive hemocytes (Fig. 5A). Fragments of other tissues that are side products of the hemocyte release method harbored EdU positive cells, serving as positive control (Fig. 5A). To explore the possibility of hemocyte proliferation upon immune challenge, we examined adults following natural infection (feeding) with the gram-negative bacterium *Serratia marcescens (S. marcescens),* or septic injury using gram-negative *E. coli, Ecc15*, or gram-positive *M. luteus*. However, even under conditions of infection we did not detect any EdU positive hemocytes (Fig. 5A). This suggests that neither differentiated macrophages, nor any potentially unlabeled progenitors giving rise to macrophages, proliferate under the tested conditions. Some hemocytes carry EdU positive cellular inclusions, which however result from phagocytosis of other EdU-incorporating polyploid or proliferative cells, rather than from hemocyte proliferation per se (Suppl.Fig. 4G-I).

**Figure 5.**
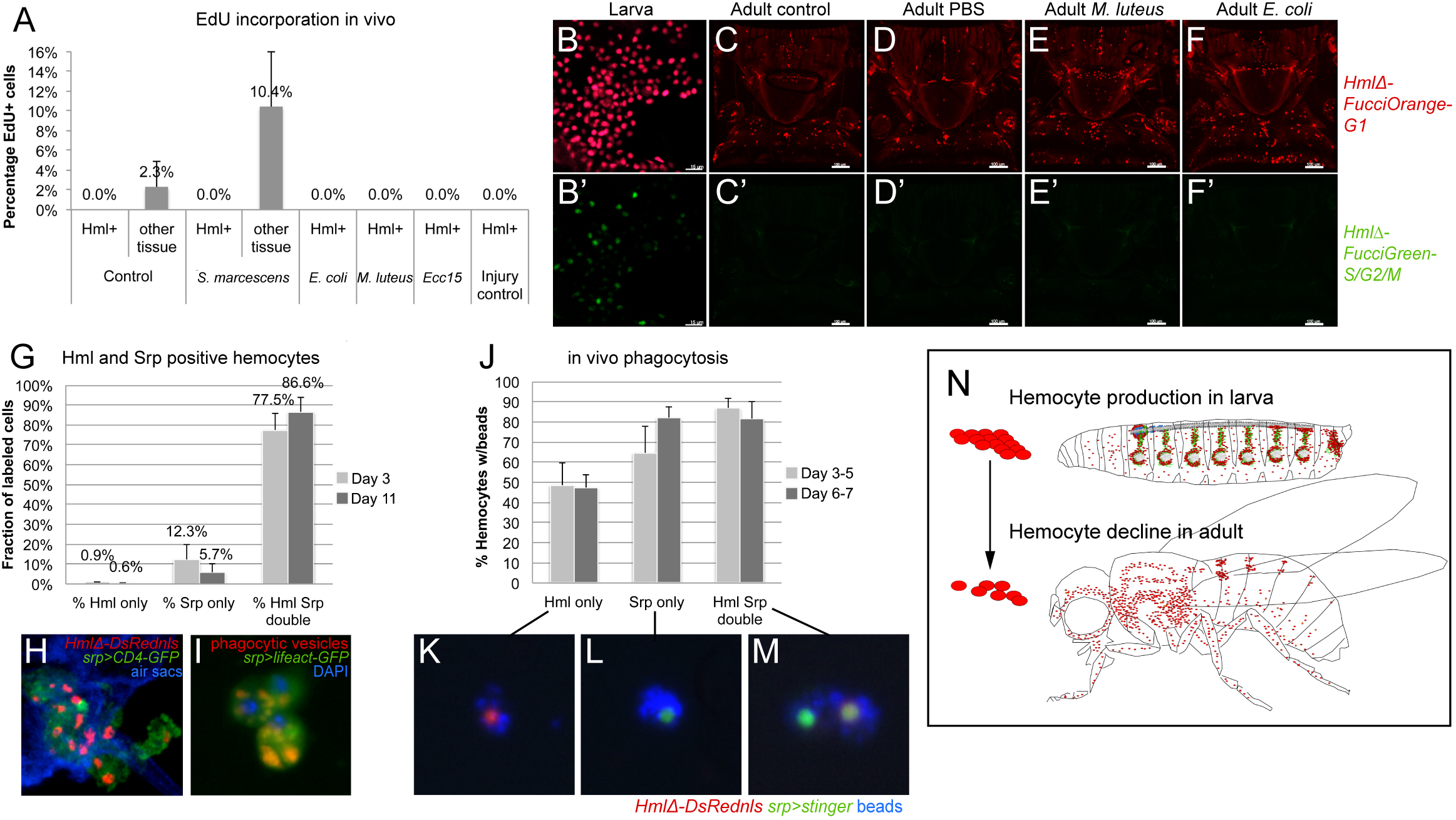
Adult hemocytes do not expand; Srp marks active phagocytes in adult *Drosophila*. In vivo EdU incorporation. Adult flies were kept on EdU containing food continuously for 2 weeks, in the absence or presence of immune challenges as indicated (see Methods). Bar chart shows percentage of EdU positive cells among hemocytes or control tissue; average and standard deviation. (B-F’) 2-color Fucci analysis of hemocytes in larvae (positive control) and adult animals, control (uninfected), sterile injury (PBS), and infection (*M. luteus, E. coli*) as indicated; genotype is *w^1118^;HmlΔFucciOrange^G1^;HmlΔFucciGreen*^*G2/S/M*^; (B-B’) embryonic-lineage hemocytes released from larvae, note presence of green cells; (C-F’) imaging of Fucci hemocytes in adult flies, dorsal views of thorax and anterior abdomen, anterior is up; note that no green hemocytes corresponding to expression of *HmlΔFucciGreen*^*G2/S/M*^ can be found. (G-I) *Srp* labels phagocytic plasmatocytes in the adult fly. (G) *Srp* and *Hml* positive hemocytes of young (3 days) and mature (11 days) adults. Note the trend of hemocytes shifting from *srp* single positive to *srp Hml* double positive with increasing maturation. (H) Thorax cross section of adult fly showing *srp* and *Hml* (double-) positive hemocytes, genotype is *HmlΔDsrednls/UAS-CD4-GFP; +/srpD-GAL4;* respiratory epithelia (air sacs, blue). (I) *Srp-Gal4, UAS-lifeact-GFP* positive plasmatocytes (green) with red phagocytic vesicles, released ex vivo from adult fly; DAPI (blue). (J-M) In vivo phagocytosis assay, based on fluorescent blue bead injection into living adult animals, followed by ex vivo examination of bead incorporation into hemocytes. (J) Quantification of hemocytes that incorporated blue beads in vivo, comparing *srp* and *Hml* single- and double positive hemocytes of young (3 days) and mature (11 days) adults. Note that *srp* single positive hemocytes show an increasing fraction of phagocytic cells upon animal maturation, and overall compare to the fraction of phagocytic hemocytes among the *srp Hml* double positives. (K, L, M) examples of labeled hemocytes as indicated. Genotype is *HmlΔDsred/ UAS-stinger; +/ srpD-GAL4.* (N) Model, hemocyte production takes place during the larval stage (mainly in the hematopoietic pockets and the lymph gland), while in the adult hemocyte numbers decline over time.

We employed numerous other methods to detect proliferation of plasmatocytes or their progenitors. Using a hemocyte-specific two-color Fucci cell cycle indicator line, we found no Fucci S/G2/M green-positive hemocytes in adult flies (Fig. 5C-F’), in contrast to proliferating larval hemocytes that served as positive control (Fig. 5B-B’). Likewise, hemocyte MARCM (Mosaic analysis with a repressable cell marker) (Lee and Luo, 1999; Makhijani et al., 2011), which labels all dividing cells that eventually would give rise to hemocytes, resulted only in minimal numbers of MARCM positive hemocytes; rare events detected could simply be due to the high background labeling seen with this assay under all induction conditions (Suppl.Fig. 4B, C) and in other systems (von Trotha et al., 2009). Lastly, PermaTwin labeling, a MARCM variant designed to detect all dividing cell progeny (Fernandez-Hernandez et al., 2013), did not produce any labeled hemocytes over induction periods of up to 3 weeks, as we determined by external imaging of intact flies and hemocyte releases. As expected, the method generated many positive cells in control tissues such as the gut (Suppl.Fig. 4D, E).

We also examined adult hemocytes for putative hemocyte progenitors. We focused on Srp, a GATA factor required for prohemocyte specification in the embryo (Rehorn et al., 1996), which was recently proposed as progenitor marker in the adult fly (Ghosh et al., 2015). We examined Srp-positive hemocytes in the adult using a Srp antibody or *srp-GAL4* driver, both of which labeled very similar, overlapping populations of hemocytes in young and more mature adults (Suppl.Fig. 4F). Surprisingly, we found that most Srp-positive hemocytes in the adult show hallmarks of phagocytosis, including internal phagocytic vesicles and the ability to phagocytose experimentally injected fluorescent microbeads (Fig. 5I-M). In maturing adults between day 3 and 11 after eclosion, an increasing number of Srp-positive hemocytes also gained expression of the plasmatocyte marker *HmlΔDsRednls* (Fig. 5G, H). However, even single positive Srp-only hemocytes are largely capable of ingesting injected beads (Fig. 5J, L), similar to *Hml* positive or double positive cells (Fig. 5K,M).

We conclude that, in the adult fly, Srp is not a marker for hemocyte progenitors. Overall, we find no indication that the adult blood cell system has significant hematopoietic capacity. Instead, we observe mainly active macrophages and a continuous decline of blood cell numbers in adult animals. Our model proposes that the adult blood cell pool results from hemocyte expansion in the larva, and over time diminishes without any significant new blood cell production (Fig. 5N).

### Hemocytes, through cell-autonomous Imd signaling, are required for an infection-induced Drosocin response

After proving that the adult blood cell population does not expand upon infection, we investigated whether the anatomical arrangement of hemocytes at the respiratory epithelia has consequences for humoral immunity. Comparing hemocyte-ablated and control animals, we examined the transcriptional induction of several antimicrobial peptide (AMP) genes following bacterial infection. Hemocyte ablation was achieved by expression of the proapoptotic genes *reaper* (*rpr*) and *head involution defect* (*hid*) (Arefin et al., 2015; Bergmann et al., 1998; Charroux and Royet, 2009; Defaye et al., 2009; White et al., 1996) and confirmed through live imaging of larvae and adults and quantification of released hemocytes (Suppl.Fig. 5A,B). *E. coli* was selected for these infections, owing to its common use as model for gram-negative infection in similar *Drosophila* studies (Ghosh et al., 2015; Lemaitre and Hoffmann, 2007). Following septic injury in the absence of hemocytes, *Drosomycin* showed little change, and *Cecropin A1* showed increased expression within the first hours of infection (Fig. 6A,B). However, expression of three AMPs was consistently reduced under hemocyte-ablated conditions, in particular *Drosocin* and, to a lesser extent, *Attacin A* and *Diptericin*, suggesting that their induction depends on hemocytes (Fig. 6C-E).

**Figure 6.**
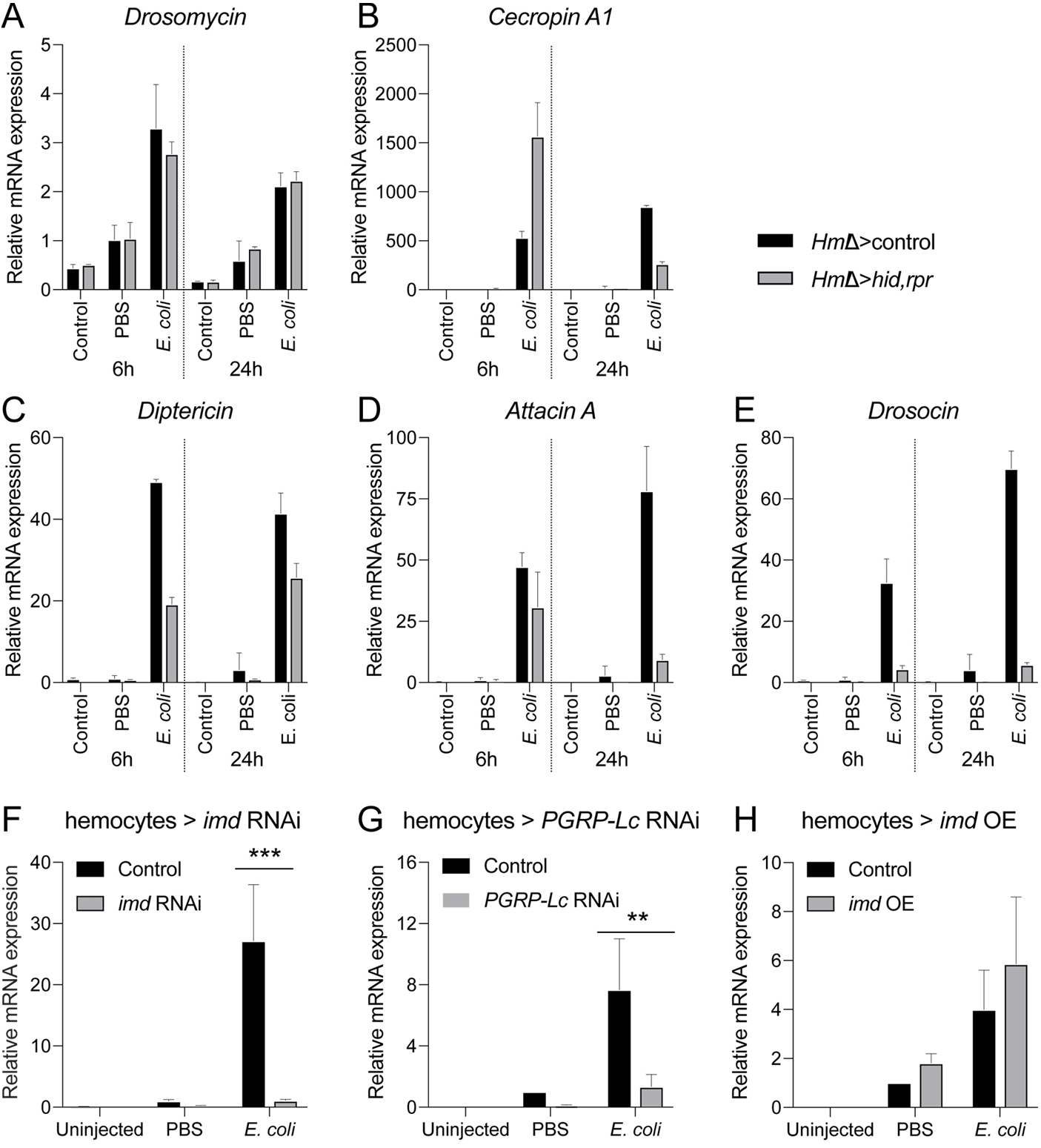
Hemocytes and Imd signaling are required for the induction of antimicrobial peptide genes including *Drosocin*. (A-E) Expression of AMPs in hemocyte-ablated flies and controls. 5 day-old adult *Drosophila* untreated, injected with sterile PBS, or with *E.coli* in PBS (OD 6), 9.2 nl; genotypes are *HmlΔ-GAL4, UAS-GFP/+* (control) or *w; HmlΔ-GAL4, UAS-GFP/ UAS-rpr; UAS-hid/+* (hemocyte ablation); flies were harvested at 6h and 24h post injection. Each chart displays mean and standard error of the mean (SEM) of samples from a representative biological replicate experiment, using pools of 10 females per condition, and triplicate qPCR runs. Values of all charts are displayed relative to the RNA level induced by the sterile PBS injections in control flies. (A) *Drosomycin*; (B) *Cecropin A1*; (C) *Diptericin*; (D) *Attacin A*; (E) *Drosocin*. (F-H) Expression of *Drosocin* in adult flies upon manipulation of Imd pathway activity. 5 day-old adult *Drosophila* untreated, injected with sterile PBS, or with *E.coli* in PBS (OD 6), 9.2 nl; flies were harvested at 6h post injection. Each chart displays the mean and confidence interval (CI) of samples from 3 averaged biological replicate experiments, using pools of 10 females per condition, and triplicate qPCR runs for each sample. Values of all charts are displayed relative to the average RNA level induced by the sterile PBS injections in control flies. Two-way ANOVA with Sidak’s multiple comparison test was performed, *,**,***,or **** corresponding to p≤0.05, 0.01, 0.001, or 0.0001, respectively (Prism). Transgenes were inducibly expressed in hemocytes 24 hours before injections. (F) *Drosocin* RNA levels of control (*HmlΔ-GAL4,UAS-GFP/+; tub-GAL80*^*ts*^*/+*) versus *HmlΔ-GAL4,UAS-GFP/+; tub-GAL80*^*ts*^*/UAS-imd RNAi*; (G) control versus *HmlΔ-GAL4,UAS-GFP/+; tub-GAL80*^*ts*^*/UAS-PGRP-LC RNAi*; (H) control versus *HmlΔ-GAL4,UAS-GFP/UAS-imd; tub-GAL80*^*ts*^*/+*.

Focusing on *Drosocin* expression as a paradigm, we asked whether TLR immune signaling in hemocytes is required for the infection-induced *Drosocin* response. We examined requirement for the NFкB-related Imd signaling pathway for detection and response to gram-negative bacteria by RNAi silencing of pathway components in hemocytes. Indeed, constitutive *imd* knockdown in hemocytes dramatically reduced *Drosocin* expression following gram-negative infection at various time points post infection (Suppl.Fig. 5C,D). Similar results were obtained when we silenced *imd* only transiently before infection (Fig. 6F), ruling out that a general shift in the immune status (Arefin et al., 2015) would cause the observed phenotype. Likewise, hemocyte-specific knockdown of the upstream peptidoglycan receptor *PGRP-LC* substantially reduced *Drosocin* induction (Fig. 6G), suggesting that recognition of bacterial cell wall components such as DAP-type peptidoglycans is required to trigger the response (Kaneko et al., 2006). Overexpression of *imd*, which leads to pathway activation (Georgel et al., 2001), mildly enhanced *Drosocin* expression after sterile and septic injury (Fig. 6H, Suppl.Fig. 5E) consistent with the Imd pathway in hemocytes being required, albeit not sufficient, for the response.

We conclude that, in response to gram-negative infection, hemocytes are essential players in the induction of humoral immunity and the expression of *Drosocin*. Signaling through the PGRP-LC/Imd pathway in hemocytes is required, albeit not sufficient, for the *Drosocin* response.

### Hemocytes are sentinels of infection that induce Drosocin expression in the respiratory epithelia and fat body

Next we examined the sites of *Drosocin* expression in the adult fly. Using a *Drosocin-GFP* transgenic reporter (Tzou et al., 2000), we found a unique expression pattern in the thorax and head of infected flies that strikingly matched locations of the hemocyte reservoir at the respiratory epithelia (Fig. 7A-C). This *Drosocin-GFP* pattern was independent of the site of bacterial injection in the thorax or abdomen of the animal (Suppl.Fig. 6A,B). We confirmed the localized *Drosocin* expression by qPCR analysis on infected flies split into two parts, one containing head and thorax, and the other consisting of abdomen only (Fig. 7D). Dissection revealed that *Drosocin-GFP* is expressed in the respiratory epithelia and in adjacent domains of the fat body, which covers the respiratory epithelia and occupies the space toward the cuticle of the fly (Fig. 7F-F’’, see also Fig. 7J-K’’’). This restriction to specific fat body domains was particularly obvious when comparing the *Drosocin-GFP* pattern to the overall expression of a fat body driver (Fig. 7E). No apparent expression of *Drosocin-GFP* was detected in hemocytes in vivo or ex vivo (Fig. 7B,F and data not shown).

**Figure 7.**
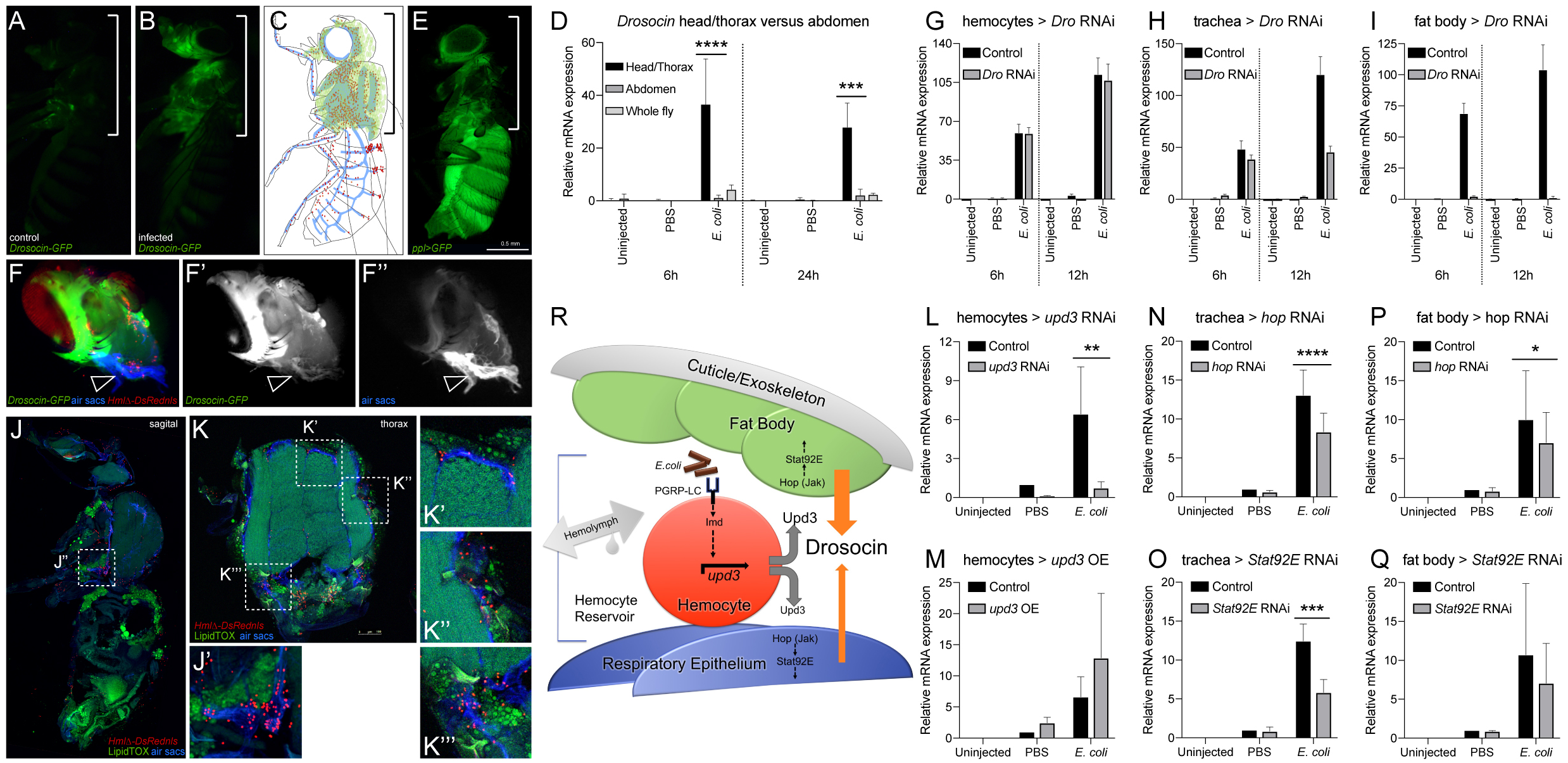
The *Drosocin* response is localized to the reservoir of hemocytes at the respiratory epithelia and colocalizing fat body domains, and requires Upd3 signaling from hemocytes. (A-C) *Drosocin-GFP* expression is restricted to the head and thorax. (A) *Dro-GFP* uninfected control; (B) *Dro-GFP* infected; (C) model of *Dro-GFP* expression (green), hemocytes (red), tracheal system (blue). (D) *Drosocin* qPCR of head/thorax versus abdomen tissue. Flies were left untreated, or injected with sterile PBS, or *E.coli* in PBS (OD 6), 9.2 nl, and harvested at 6 and 24h post infection. Two-way ANOVA with Sidak’s multiple comparison test was performed for head/thorax versus abdomen, *,**,***,or **** corresponding to p≤ 0.05, 0.01, 0.001, or 0.0001, respectively. (E) Location of fat body throughout the animal marked by *ppl-GAL4, UAS-GFP* (green). (F-F’’) Dissected heads of genotype *Drosocin-GFP/HmlΔ-DsRed* (Drosocin-GFP green, hemocytes red), respiratory epithelia (air sacs, blue). Note *Drosocin-GFP* expression is high in fat body and moderate in respiratory epithelia (arrowhead). (G-I) Tissue specific RNAi knockdown of *Drosocin*; overall *Drosocin* mRNA levels were quantified by qPCR. 6-7 day-old adult females were left untreated, insejcted with PBS, or *E.coli* in PBS (OD 6), 9.2 nl, and harvested 6 and 12h post infection. Each chart displays mean and SEM of samples from a representative biological replicate experiment, using pools of 10 females per condition, and triplicate qPCR runs. Values of all charts are displayed relative to the RNA level induced by the sterile PBS injections in control flies. (G) *Drosocin* RNAi silencing in hemocytes; (H) in respiratory system; (I) in fat body. (J-K’’’) Anatomy of fat body tissue lining the respiratory epithelia and hemocytes; *HmlΔ-DsRednls* (hemocytes, red), fat body (LipidTOX, large, distinct green cells), respiratory epithelia (air sacs, blue) (J) Sagital section of adult *Drosophila*. (J’) Closeup of region indicated in (J). (K’-K’’’) Closeup of regions indicated in (K). Note that hemocytes are layered between tracheal tissue and fat body. (L-Q) Expression of *Drosocin* in adult flies upon silencing or overexpression of *upd3* and silencing of genes of the Jak/Stat pathway. 5 day-old adult *Drosophila* untreated, injected with sterile PBS, or *E.coli* in PBS (OD 6), 9.2 nl; flies were harvested at 6h post injection. Each chart displays the mean and CI of samples from 3 averaged biological replicate experiments, using pools of 10 females per condition, and triplicate qPCR runs for each sample. Values of all charts are displayed relative to the average RNA level induced by the sterile PBS injections in control flies. Two-way ANOVA with Sidak’s multiple comparison test was performed, *,**,***,or **** corresponding to p≤0.05, 0.01, 0.001, or 0.0001, respectively. (L, M) *Drosocin* qPCR of whole flies, inducible transgene expression in hemocytes, (L) Genotypes are control (*HmlΔ-GAL4,UAS-GFP/+; tub-GAL80*^*ts*^ */+*) versus *HmlΔ-GAL4,UAS-GFP/+; tub-GAL80*^*ts*^ */UAS-upd3* RNAi. (M) Control versus *HmlΔ-GAL4,UAS-GFP/ UAS-upd3; tub-GAL80*^*ts*^ */+.* (N, O) *Drosocin* qPCR of whole flies, inducible transgene expression in tracheal system. (N) Genotypes are control *(btl-GAL4, tub-GAL80*^*ts*^, *UAS-GFP / +*) versus *btl-GAL4, tub-GAL80*^*ts*^, *UAS-GFP / UAS-hop* RNAi; (O) Genotypes are control versus *btl-GAL4, tub-GAL80*^*ts*^, *UAS-GFP / UAS-Stat92E* RNAi. (P, Q) *Drosocin* qPCR of whole flies, transgene expression in fat body. (P) Genotypes are control (*ppl-GAL4, UAS-GFP* / +) versus *ppl-GAL4, UAS-GFP* /+; *UAS-hop* RNAi/+; (Q) Genotypes are control versus *ppl-GAL4, UAS-GFP*/+; *UAS-Stat92E* RNAi/+. (R) Model of communication between hemocytes, fat body and respiratory epithelia, in which hemocytes act as sentinels of infection. Gram-negative bacteria that accumulate together with hemocytes in the reservoir between respiratory epithelia and fat body trigger activation of the Imd pathway through the peptidoglycan recognition protein PGRP-LC in hemocytes. Imd signaling drives induction of *upd3* expression in hemocytes, leading to Upd3 secretion. Upd3 activates Jak/Stat signaling in adjacent domains of the fat body and the respiratory epithelia, contributing directly or indirectly to the induction of *Drosocin* expression.

In order to further confirm the contribution of each tissue to the overall *Drosocin* response, we performed tissue specific RNAi knockdowns of *Drosocin* and analyzed the levels of total *Drosocin* expression by qPCR of whole flies. Silencing of Drosocin in the respiratory system shows partial contribution to the total *Drosocin* expression (Fig. 7H). Silencing of *Drosocin* in fat body demonstrated that the major contribution comes from this tissue (Fig. 7I), consistent with the observed high levels of *Drosocin-GFP* expression in fat body, and its established role as major site of AMP expression. In contrast, *Drosocin* silencing in hemocytes caused little to no reduction in overall *Drosocin* levels, again confirming that hemocytes themselves are not a significant source of total *Drosocin* expression in the fly (Fig. 7G). Overall *Drosocin* knockdown efficiency was close to 100% using a ubiquitous driver (Suppl.Fig. 6D).

Considering that hemocytes are required for *Drosocin* induction, yet *Drosocin* is predominantly expressed in domains of the fat body and the respiratory epithelia, we hypothesized that hemocytes might be required as sentinels of infection that signal through some molecular mechanism to the *Drosocin*-expressing tissues. Cryosections of whole adult animals and head dissections show that hemocytes tightly colocalize with, and are layered in between, the respiratory epithelia and fat body, suggesting an interface that facilitates signaling communications among the tissues (Fig. 7J-K’’’, Suppl.Fig. 6C,C’). Hypothesizing communication via a secreted factor, we tested potential candidate genes of secreted signaling molecules by RNAi and overexpression. One gene that stood out was *unpaired 3* (*upd3* (Agaisse et al., 2003)), a ligand of the Jak/Stat pathway in *Drosophila*. Hemocyte specific knockdown of *upd3* strongly reduced *Drosocin* expression after gram-negative infection (Fig. 7L), suggesting that Upd3 is a required hemocyte produced signal in the communication to the respiratory epithelia and fat body. Conversely, *upd3* overexpression mildly enhanced the response under conditions of sterile and septic injury (Fig. 7M), resembling the effect of *imd* overexpression (see Fig. 6H) and suggesting that *upd3* in hemocytes is required albeit not sufficient.

Since Upd3 is a ligand of the Jak/Stat pathway, we also probed for the requirement of pathway components in the putatively receiving tissues. Indeed, RNAi silencing of the pathway components *hopscotch* (*hop*) (*Drosophila* Jak) or *Stat92E* in the respiratory system partially reduced the overall *Drosocin* response (Fig. 7N, O). Likewise, silencing of *hop* or *Stat92E* in the fat body also led to a partial reduction in total *Drosocin* expression (Fig. 7P,Q). This suggests that Jak/Stat signaling in both the respiratory epithelia and fat body are required to respond to hemocyte-expressed Upd3 and to contribute jointly to overall *Drosocin* expression, consistent with our earlier findings. Unexpectedly, overexpression of activated Jak *hop*^*TumL*^ in fat body, or activated *Stat92E* (combined *Stat92E; Stat92E*^*ΔNΔC*^) in the respiratory system or fat body, reduced *Drosocin* levels after infection (Suppl. Fig. 6E-G). Crosses of *UAS-hop*^*TumL*^ with the tracheal driver *btl-GAL4* were largely lethal and gave rise to only a minimal number of escapers, despite the use of *tub-GAL80*^*ts*^ to avoid expression during development. The unexpected effects of Jak/Stat overexpression might be due to some unexpected activation of a negative feedback loop or other complex signaling changes.

Taken together, our findings suggest that hemocytes act as sentinels of bacterial infection, which signal to neighboring cells of the respiratory epithelia and fat body. Hemocyte-expressed *upd3* and Jak/Stat signaling in the respiratory epithelia and fat body are required, albeit not sufficient, in this process. In response, both fat body and respiratory epithelia concertedly contribute to the expression of *Drosocin*.

### Drosocin expression in the respiratory epithelia and fat body promote animal survival after infection

Lastly, we asked whether *Drosocin* expression in the respiratory epithelia and fat body is crucial for animal survival after infection. First we determined whether ubiquitous RNAi silencing of *Drosocin* affects animal survival after immune challenge. Indeed we found that survival after bacterial infection with gram-negative *E. coli* or *E. cloacae* was significantly reduced, an effect seen with two independent *Drosocin* RNAi lines (Fig. 8A,D). *Drosocin* knockdown also affected survival after injury (PBS injection) (Fig. 8B). Ubiquitous knockdown of *Drosocin* did not affect the induction of other AMPs following bacterial infection (Suppl.Fig. 7A-D), strengthening the idea that endogenous Drosocin has direct antibacterial role/s, and does not act as signal or mediator that would sustain the expression of other AMPs following bacterial infection.

**Figure 8.**
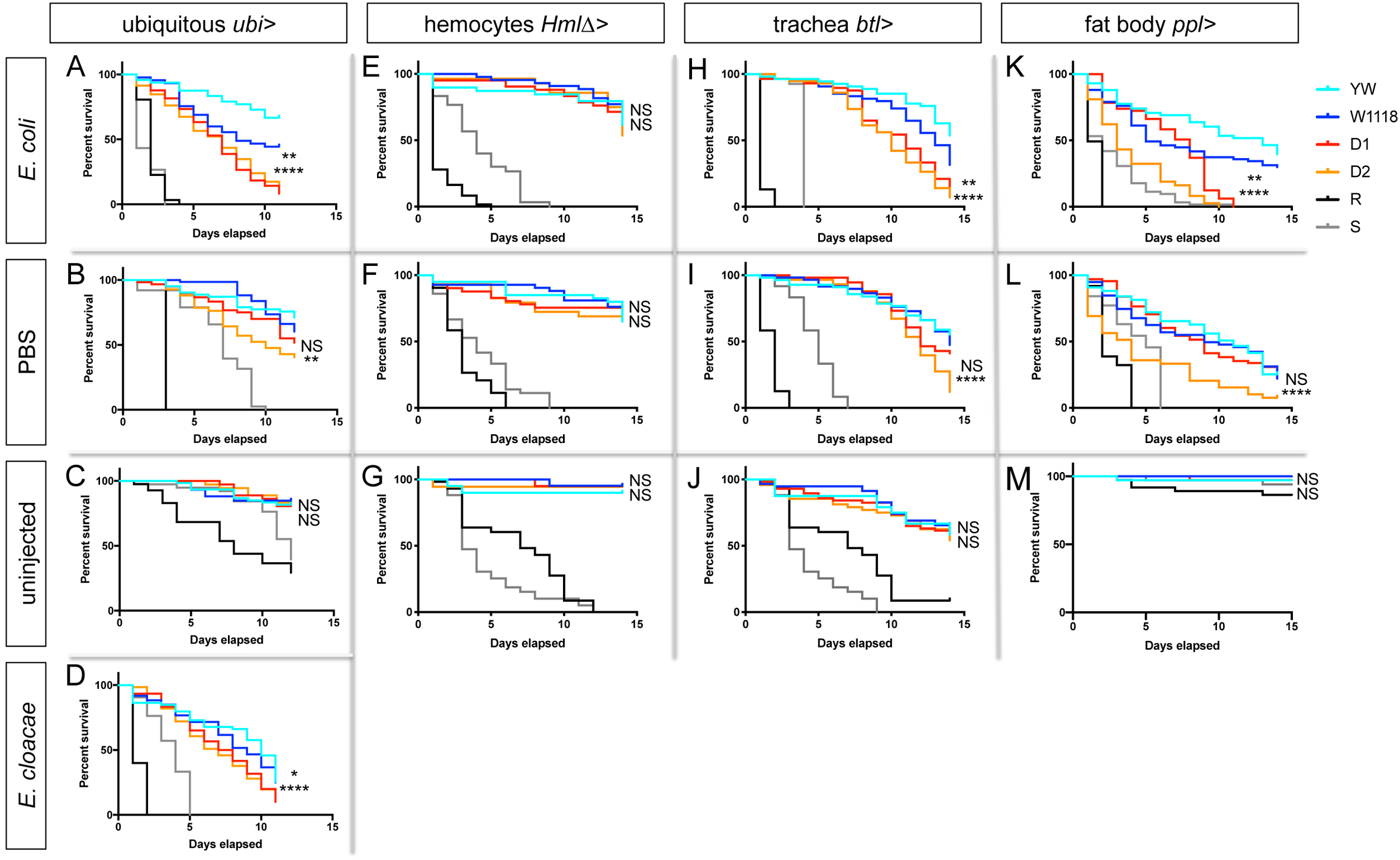
*Drosocin* silencing in respiratory epithelia and fat body decreases animal survival after infection. (A-D) Survival assays of adult female F1 progeny resulting from crosses of GAL4 drivers with the following transgenic lines were analyzed: *UAS-Drosocin RNAi* lines (D1, D2) or controls (*yw* (YW), *w^1118^* (W1118)). Mutant strains *Rel*^*E20*^ (R) and *spz*^*rm7*^ (S) served as controls. Figure displays one out of 3 comparable biological replicate experiments; in each experiment, for each genotype and condition 40 to 60 females were assessed; p-values log-rank (Mantel-Cox) test, *,**,***,or **** corresponding to p≤0.05, 0.01, 0.001, or 0.0001, respectively; upper symbol corresponds to comparison *w1118* versus *Dro* RNAi D1; lower symbol corresponds to comparison of crosses of *yw* versus *Dro* RNAi D2. 5 day old female flies were treated as follows and then incubated at 29°C. (A-D) *Ubi-GAL4* crosses, (A) *E. coli* (OD6, 9.2 nl); (B) sterile PBS injection (9.2nl); (C) uninjected control; (D) *E. cloacae* (OD4, 9.2nl). Note that *Drosocin* knockdown caused significantly reduced survival after gram-negative infection. After PBS injection, survival was significantly reduced for D1 in 2 out of 3 replicate experiments, for D2 in 3 out of 3 replicate experiments. (E-G) *HmlΔ-GAL4* crosses, (E) *E. coli* (OD6, 9.2 nl); (F) sterile PBS injection (9.2nl); (G) uninjected control. (H-J) *btl-GAL4* crosses, (H) *E. coli* (OD6, 9.2 nl); (I) sterile PBS injection (9.2nl); (J) uninjected control. (K-M) *ppl-GAL4* crosses, (K) *E. coli* (OD6, 9.2 nl); (L) sterile PBS injection (9.2nl); (M) uninjected control. Note that ubiquitous *Drosocin* knockdown caused significantly reduced survival after gram-negative infection (A, D). After PBS injection, survival was significantly reduced (B), for D1 in 2 out of 3 replicate experiments, for D2 in 3 out of 3 replicate experiments. *Drosocin* knockdown in tracheal system or fat body caused significantly reduced survival after gram-negative infection (H, K). *Drosocin* knockdown in the tracheal system or fat body showed a partially penetrant effect on survival after PBS injection (I, L); in tracheal system knockdown, survival was significantly reduced for D1 in 0 out of 3 replicate experiments, for D2 in 2 out of 3 replicate experiments. For fat body knockdown, survival was significantly reduced for D1 in 1 out of 3 replicate experiments, for D2 in 1 out of 3 replicate experiments. As expected, survival is not affected when *Drosocin* is silenced in hemocytes (E, F, G).

Next we probed the role of *Drosocin* expression specifically in the respiratory system or fat body. Indeed, *Drosocin* silencing in the respiratory system or fat body significantly reduced survival after *E. coli* infection (Fig. 8H,K), an effect that was again seen with two independent *Drosocin* RNAi lines. *Drosocin* expression in the respiratory system and fat body may also contribute to survival after injury, as *Drosocin* knockdown in either of the two tissues showed a partially penetrant effect on survival after PBS injection (Fig. 8I,L). As expected, silencing of *Drosocin* in hemocytes had no significant effect on survival following gram-negative infection or injury (Fig. 8E,F).

Taken together, we conclude that endogenous *Drosocin* expression in the respiratory system and fat body of adult *Drosophila* is effective in promoting animal survival after gram-negative infection. This protective role of localized *Drosocin* expression is directly linked to the reservoirs of blood cells in adult *Drosophila*. Our findings highlight the role of hemocytes as sentinels of infection that relay the innate immune response to colocalizing domains of the fat body and the respiratory epithelia. By identifying Imd signaling and upd3 expression in hemocytes as the first molecular steps in this relay, our work establishes a new *Drosophila* model to study multi-tissue organismal immunity.

## Discussion

Adult *Drosophila* has been a powerful system to dissect molecular mechanisms underlying innate immunity, from pattern recognition, TLR- and cytokine-related signaling cascades, to humoral effectors such as antimicrobial peptides (Imler and Bulet, 2005; Lemaitre and Hoffmann, 2007; Morin-Poulard et al., 2013; Royet and Dziarski, 2007) However, major questions have remained regarding the hematopoietic capacity of the *Drosophila* adult blood cell system, and the role of blood cells in organismal immunity. Addressing these questions, we discovered a central role for a new blood cell reservoir at the respiratory epithelia and fat body of adult *Drosophila*. The reservoir serves as major receptacle of blood cells and foreign particles, and in addition executes a local humoral immune response of Drosocin expression that promotes animal survival after bacterial infection. Both functions are tied together by hemocytes acting as sentinels of infection, that signal through the Imd pathway and Upd3 to induce the *Drosocin* in the tissues of their surrounding reservoir, i.e. the respiratpry epithelia and colocalizing domains of the fat body.

In the past, studies on *Drosophila* and other insects typically focused on the adult heart as the site of hemocyte accumulation. Historic insect literature described clusters of hemocytes at the ostia of the heart as “immune organ” (Gupta, 2009), which were confirmed as locations where hemocytes and bacteria accumulate (King and Hillyer, 2012). More recently, a model of adult blood cell production at the heart was proposed (Ghosh et al., 2015). Some studies described functions of hemocytes in other locations, such as at the ovaries or along the gut of adult flies (Ayyaz et al., 2015; Van De Bor et al., 2015). In our study we took a more global approach. Surveying the whole animal by cryosectioning afforded us to identify the largest reservoir of hemocytes in adult *Drosophila*, which is surrounding the respiratory epithelia of the thorax and head. The respiratory hemocyte reservoir is lined by fat body, a major immune tissue, and is directly connected with the open circulatory system (Fig. 7R). Our evidence from microbead- and bioparticle injections and developmental studies suggest that hemocytes and particles are delivered to these areas by the streaming hemolymph, even though the detailed anatomy of the open circulatory system remains to be mapped in more detail. Hemocytes may get physically caught in these locations, or in addition may use a more active adhesion mechanism., The intimate relationship of hemocytes with the respiratory epithelia, hemolymph, and adjacent fact body may serve dual interconnected roles, (1) guarding the respiratory epithelia as a barrier to the environment with respect to the roles of hemocytes in both phagocytosis and the induction of humoral immunity, and (2) facilitating gas exchange of hemocytes and nearby immune tissues, which in turn may again benefit defense functions. The former may be particularly advantageous in the defense against fungal pathogens, such as the entomopathogenic fungus *B. bassiana*, which invades *Drosophila* via the tracheal system as primary route of infection (Clarkson and Charnley, 1996; Lemaitre et al., 1997). Regarding the latter, a study in caterpillars described the association of hemocytes with trachea, proposing a function for the respiratory system to supply hemocytes with oxygen (Locke, 1997).

*Drosophila* adult blood cells derive from two lineages: one that originates in the embryo and resembles vertebrate tissue macrophages, and another that produces blood cells in the lymph gland through a progenitor-based mechanism (Banerjee et al., 2019; Gold and Brückner, 2014, 2015; Holz et al., 2003) Based on flipout-*lacZ* lineage tracing we estimate that more than 60% of adult hemocytes derive from the embryonic lineage, inferring that the rest derive from the lymph gland. This is surprising, considering older reports that proposed the majority of adult hemocytes would derive from the lymph gland (Lanot et al., 2001), Our findings places more importance on the *Drosophila* embryonic lineage of hemocytes and suggest additional parallels with tissue macrophages in vertebrates, which persist into adulthood and form a separate myeloid system independent of the progenitor-derived monocyte lineage (Davies et al., 2013; Gold and Brückner, 2014, 2015; Perdiguero et al., 2014; Sieweke and Allen, 2013). Future research will show whether the relative contribution of the two hemocyte lineages to the adult blood cell pool will be the same or different under conditions of stress and immune challenges.

Given that embryonic-lineage plasmatocytes are highly proliferative in the hematopoietic pockets of the *Drosophila* larva, and lymph gland hemocyte progenitors and some lymph gland plasmatocytes proliferate during larval development (Evans et al., 2003; Jung et al., 2005; Letourneau et al., 2016), the absence of hemocyte proliferation in the adult is surprising. However, no significant new plasmatocyte production, neither from direct proliferation nor a proliferating progenitor, was detected using a variety of approaches including 2-week in vivo EdU incorporation, Fucci, PermaTwin- and hemocyte MARCM analyses, all under standard laboratory conditions and bacterial infection.

Two additional lines of evidence support a lack of hematopoietic activity in adult *Drosophila*. First, quantifications of total hemocytes ex vivo demonstrate a continuous decline of blood cells in the animal over time, even under conditions of immune challenge. Second, we find no evidence that Srp in adult *Drosophila* is a progenitor marker, as Ghosh et al. postulated in support of their adult hematopoiesis model (Ghosh et al., 2015). Instead, we demonstrate that the vast majority of Srp-expressing hemocytes in the adult are active phagocytes, therefore lacking hallmarks of undifferentiated progenitors. This contrasts the above model (Ghosh et al., 2015), and reveals differences to embryonic development, where Srp is required for the specification of undifferentiated prohemocytes (Rehorn et al., 1996)

As an advantage to our study, we globally examined *Drosophila* adults for new blood cell production, taking into account all hemocyte populations of the animal. This allowed us to clarify a process that we believe formed another pillar in the model claiming blood cell production at the heart of adult *Drosophila* (Ghosh et al., 2015). During maturation of the adult animal, hemocytes relocate to the respiratory epithelia and heart upon completion of cytolysis of larval fat body cells (Nelliot et al., 2006). This recruitment increases local hemocytes visible at the heart and in areas underlying the cuticle, but does not increase the absolute number of blood cells. However, when focusing on this area in isolation, the false impression of an expansion in the blood cell population could arise. Similarly, we do not discern a significant increase in total blood cells following bacterial infection, although increased numbers of fluorescently labeled hemocytes become visible through the cuticle. We attribute this to the infection-induced upregulation of hemocyte-specific genes such as *Hml* and *crq* and their respective enhancers in e.g. *HmlΔ-GAL4*, driving expression of *UAS-GFP*. Enhanced hemocyte expression of these genes post infection was also described previously (De Gregorio et al., 2001; Franc et al., 1999).

Taken together, our broad evidence speaks to a lack of significant hematopoietic capacity of the blood cell system in adult *Drosophila*. Our findings are in agreement with other studies that have reported a lack of hemocyte proliferation in adult *Drosophila* (Lanot et al., 2001; Mackenzie et al., 2011; Woodcock et al., 2015), and functional immunosenescence in ageing flies (Felix et al., 2012; Mackenzie et al., 2011). Of course we cannot exclude the possibility that some other specific immune challenge or stress might exist that would be potent enough to trigger proliferation- or differentiation-based blood cell production in adult *Drosophila*. Likewise, although the majority of Srp positive hemocytes are active macrophages, we cannot exclude that adult *Drosophila* may possess small numbers of proliferation-competent progenitors that may have persisted e.g. from the lymph gland posterior lobes (Grigorian et al., 2011; Jung et al., 2005); such cells might give rise to new differentiated hemocytes, although according to our data they would remain insignificant in number.

Taking into account the relatively short life span and short reproductive phase of *Drosophila*, the adult fly may be sufficiently equipped with the pool of hemocytes that is produced in the embryo and larva. In fact, hemocytes may not be essential for the immediate survival of adult flies (see below). We propose a model that places emphasis on larval development as the sensitive phase for the expansion and regulation of the adult blood cell pool (Gold and Brückner, 2015) (Fig. 5N). In the larva, both the embryonic-lineage hemocytes in the hematopoietic pockets and blood cells deriving from the lymph gland integrate signals from a variety of internal and external stimuli to adapt to existing life conditions (Gold and Brückner, 2014, 2015; Shim, 2015).

*Drosophila* ablated of hemocytes, and mutants devoid of hemocytes, survive to adulthood (Arefin et al., 2015; Braun et al., 1998; Charroux and Royet, 2009; Defaye et al., 2009). However, hemocyte-depleted animals have been known to be more prone to, and succumb more rapidly to, infection (Arefin et al., 2015; Basset et al., 2000; Braun et al., 1998; Charroux and Royet, 2009; Defaye et al., 2009). Our work reveals a role for hemocytes in a local humoral immune response of the fat body and respiratory epithelia. Previous studies on hemocyte-ablated flies have reported increases in *Defensin* and *IM1* expression (Charroux and Royet, 2009; Defaye et al., 2009). Likewise, we find that hemocyte ablation enhances initial *Cecropin A1* expression after infection. This may indicate a negative regulatory role of hemocytes on these and possibly other AMP genes. It may also be directly or indirectly linked to the altered inflammatory status of hemocyte-ablated flies, which show increased Toll and decreased Imd signaling (Arefin et al., 2015). In contrast, we find a positive role for hemocytes in the induction of *Attacin A, Diptericin, and Drosocin*. Expression of *Drosocin* is particularly intriguing, as its pattern of expression matches the tissues that form the hemocyte reservoir, i.e. the respiratory epithelia and fat body domains of the head and thorax. The concept of hemocytes promoting AMP expression in other tissues is well established (Agaisse et al., 2003; Basset et al., 2000; Chakrabarti et al., 2016; Yang et al., 2015). A role for AMP expression in surface epithelia that interface with the environment was reported by Ferrandon et al. (Ferrandon et al., 1998), and *Drosocin* expression was described in embryonic and larval trachea and the abdominal tracheal trunks of adult *Drosophila*, albeit not in the respiratory epithelia (Akhouayri et al., 2011; Tan et al., 2014; Tzou et al., 2000). *Attacin A* is also expressed in the larval trachea (Akhouayri et al., 2011), warranting future investigation into its expression in the adult respiratory epithelia and putative regulatory links with hemocytes. In adult *Drosophila*, hemocytes tightly localize between the respiratory epithelia and a layer of adult fat body tissue that occupies the space toward to the cuticle exoskeleton, forming an anatomical and functional triad of hemocytes, respiratory epithelia and fat body, in which hemocytes act as sentinels of infection.

We propose that this close anatomical relationship facilitates rapid local signaling by (1) detection of gram-negative bacteria through PGRP-LC/Imd signaling in hemocytes and (2) communication of hemocytes with the fat body and respiratory epithelia through Upd3/Jak/Stat signaling, culminating in the induction of *Drosocin* expression in these tissues to protect the animal following bacterial infection (Fig. 7R). Consistent with previous knowledge that *Drosocin* expression is lost in *imd* mutant backgrounds (Lemaitre PNAS 1995; Rahel et al. JBC 2004), we find that hemocyte-autonomous Imd signaling is required, albeit not sufficient, to trigger the infection-induced *Drosocin* response. Likewise, the Imd pathway upstream receptor *PGRP-LC* is required in hemocytes, suggesting that DAP-type peptidoglycan recognition and initiation of Imd signaling are a critical step in triggering the *Drosocin* response. Our data suggest roles for hemocyte-expressed *upd3*, and corresponding Jak/Stat signaling in cells of the fat body and respiratory system, all of which are required albeit not sufficient. Overexpression of activated *Stat92E* (combined *Stat92E; Stat92E*^*ΔNΔC*^) or the activated Jak *Hop*^*TumL*^ paradoxically suppress *Drosocin* expression. Crosses of *UAS-hop*^*TumL*^ with the tracheal driver *btl-GAL4* were largely lethal despite our use of *tub-GAL80*^*ts*^, which might indicate leaky expression of the *UAS-hop*^*TumL*^ transgene. Overall, we can only speculate that the unexpected effects of Jak/Stat overexpression might be due to activation of some negative feedback loop or other complex signaling changes.

Several reports provide precedent for a role of hemocyte-expressed Upd3 in the induction of immune responses in other target tissues. Following septic injury, upregulation of *upd3* in hemocytes triggers induction of stress peptide genes of the *turandot* family including *totA* in fat body (Agaisse et al., 2003; Brun et al., 2006). Similarly, in response to injury, hemocyte-produced Upd3 induces Jak/Stat signaling in the fat body and gut (Chakrabarti et al., 2016). Further, under lipid-rich diet, *upd3* is induced in hemocytes, causing impaired glucose homeostasis and reduced lifespan in adult *Drosophila* (Woodcock et al., 2015). Lastly, in the larva, hemocyte-derived Upd2 and −3 activate Jak/Stat signaling in muscle, which are required for the immune response against parasitic wasps (Yang et al., 2015). However, in the *Drosocin* response around the reservoir of hemocytes, our data predict that additional signal/s and/or signaling pathway/s are needed to initiate *Drosocin* expression and potentially restrict its expression to defined fat body domains of the head and thorax. Additional events may rely on diverse mechanisms. They could include signaling through Toll or other signaling pathways in hemocytes and/or other tissues including the respiratory epithelia and fat body. Likewise, other types of signals may be required or permissive for the *Drosocin* response in the adult fly, such as reactive oxygen species (ROS) or nitric oxide (NO) which were reported to play roles in the relay of innate immune responses to infection and stress (Caceres et al., 2011; Di Cara et al., 2018; Eleftherianos et al.,2014; Myers et al., 2018; Wu et al., 2012), or non-peptide hormones including ecdysone, which confers competence in the embryonic tracheal *Drosocin* response to bacterial infection, and enhances humoral immunity under conditions of dehydration (Tan et al., 2014; Zheng et al., 2018). Lastly, there could be e.g. a requirement for additional processing to make bacterial ligands accessible for receptors in other tissues, as has been reported for Psidin, a lysosomal protein required in blood cells for degradation of engulfed bacteria and expression of *Defensin* in the fat body (Brennan et al., 2007), although this mechanism may not be universal in all systems (Nehme et al., 2007).

Our work reveals an active role of endogenous *Drosocin* expression in survival after bacterial infection. Since the cloning of *Drosocin* and its classification as an inducible antibacterial peptide (Bulet et al., 1993), *Drosocin* has been studied for its transcriptional regulation (Charlet et al., 1996), illustrating its induction under a variety of bacterial and other immune challenges (Akbar et al., 2011; Akhouayri et al., 2011; Becker et al., 2010; Clark et al., 2013; Fernando et al., 2014; Gendrin et al., 2013; Lemaitre et al., 2012; Yagi et al., 2013). Drosocin structure and antimicrobial function have been studied in vitro (Ahn et al., 2011; Bikker et al., 2006; McManus et al., 1999; Otvos et al., 2000b), and by overexpression from transgenes in *Drosophila* (Loch et al., 2017; Tzou et al., 2002; Vonkavaara et al., 2013) and in heterologous vertebrate systems (Otvos et al., 2000a). Consistent with our findings, a recent study that examined new CRISPR-based *Drosocin* null mutants reached similar conclusions regarding the requirement of endogenous *Drosocin* expression for animal survival following *E. cloacae* infection (Hanson et al., 2019). However, this study did not address anatomical features of *Drosocin* expression, nor its unique path of induction. In addition to *Drosocin*’s role in animal survival after bacterial infection, our data suggest contribution of *Drosocin* to animal survival after injury (PBS injection); however, due to the possibility of inadvertent contamination of the injection site we cannot exclude that this effect may also involve aspects of infection. A role for endogenous Drosocin levels in the antimicrobial response is strongly supported by independent data in the literature. Specifically, the minimum inhibitory concentration (MIC) of Drosocin against *E. coli* and *E. cloacae* was determined to be well within the range or below the endogenous concentration of Drosocin in the *Drosophila* hemolymph (MIC is 1 µM or 2µM for the glycosylated forms, and 8 or 10µM for the unglycosylated form, respectively (Bulet et al., 1996), compared to 40µM Drosocin in the *Drosophila* hemolymph ((Uttenweiler-Joseph et al., 1998)).

In conclusion, our work recognizes adult *Drosophila* as a promising model to study organismal immunity centering around the reservoir of blood cells, which involves immune signaling in the triad of hemocytes, respiratory epithelia and fat body. At the evolutionary level, this model shows parallels with vertebrate immune cells of the lung and innate immune responses to bacterial infection (Byrne et al., 2015; Divangahi et al., 2015; Opitz et al., 2010). The *Drosophila* model opens countless new avenues for exciting future research, for example to investigate additional molecular and cellular mechanisms in the immune signaling relay, the role and regulation of the system in the defense against pathogens that invade the trachea as natural route of infection, the anatomical and functional features of the respiratory epithelia and other parts of the hemocyte reservoir, features of the open circulatory system and its particle streaming dynamics, effects of the blood cell reservoir on hemocyte activity, the impact of other tissues and systemic factors on the immune response, the use of the same axis by gram-positive or non-bacterial pathogens, and the induction of other AMPs and immune effector genes in the same axis of regulation.

## Experimental Procedures

### *Drosophila* strains and fly husbandry; *HmlΔFucci* transgenic lines; Flipout-*LacZ* lineage tracing; Fucci analysis and quantification of hemocytes; Hemocyte MARCM; PermaTwin MARCM; Quantification of AMP expression by qPCR

Please see Supplemental Experimental Procedures.

#### Cryosectioning, head dissections

For cryosectioning, adult whole flies were embedded in OCT (Tissue-Tek) and snap frozen on dry ice with 95% ethanol. Cryosections of 55µM thickness were obtained using a cryostat (Leica CM3050 S or Leica CM1950). Tissue sections were placed on charged glass slides and fixed with 4% PFA in 2xPBS with 2x Complete protease inhibitor (Roche) for 10 min, followed by 2% PFA for 10 min, and permeabilized with 0.2% Triton-X100 for 15 min. Dissections of the respiratory epithelia (air sacs) of the head were done using forceps and a dissection well filled with PBS. Heads were detached from the rest of the body. By holding heads at the proboscis, the cuticle of the head was opened through the antennae first, progressively working toward the posterior, trying to avoid damage of the respiratory epithelia. Once the brains with the overlying respiratory air sacs were exposed, they were fixed with 4% PFA/PBS for 10 minutes, washed in PBS and processed for imaging. In both cryosections and dissections, hemocyte attachment to the respiratory epithelia is quite fragile. Accordingly, some hemocyte loss occurs in many preparations.

### Immunohistochemistry and other staining methods

To analyze hemocytes ex vivo, hemocytes of single flies were released in 100μl of Schneider’s medium (Gibco, Millipore) or PBS supplemented with Complete 2x (Roche) and proteinase inhibitor 4-(2-Aminoethyl)benzenesulfonyl fluoride hydrochloride AEBSF (Sigma); flies were dissected and hemocytes were scraped from all inside areas of the fly (head, thorax and abdomen).

For immunohistochemistry, antibodies used were goat anti-GFP (Rockland Immunochemicals, 1:2000), rabbit anti-βGal (Thermo, 1:1000), rabbit anti-DsRed (Rockland Immunochemicals, 1:1000), mouse P1 (P1a+P1b) (Kurucz et al., 2007) (kind gift of I. Ando, 1:10), anti-Crq (Franc et al., 1996) (kind gift of C. Kocks, 1:1000), anti-Srp (kind gift of A. Giangrande, 1:1000) and secondary antibodies conjugated to Alexa dyes (Molecular Probes, 1:500), fluorescently labeled phalloidin (Molecular Probes), DAPI (Sigma), DRAQ5 (ThermoFisher).

Fat body cells were labeled using OilRedO (37%) dissolved in triethyl phosphate (6ml triethyl phosphate and 4 ml water) for 30 min, followed by three to four washes with distilled water. Other stainings used fluorescent LipidTOX dyes (LifeTech), diluted in PBS.

### Edu incorporation assays

To assess cell proliferation in adult hemocytes, adult F1 progeny of *yw* or *HmlΔ-GAL4, UAS-GFP x CantonS* were maintained for 14 days on fly food supplemented with a stock of 0.5 mg/ml EdU (Invitrogen) to a final concentration of 0.4mM EdU. Animals were immune challenged with bacterial infections both before or during EdU feedings. Various ages of adult animals (starting labeling at 0-1 week after eclosion) and crosses with other control lines (Oregon R, w1118 etc.) were examined. Hemocytes were released in multi-well dishes ex vivo and Click-IT EdU detection was performed according to the manufacturer’s instructions (Invitrogen).

### Microbead and bioparticle injections

For microbead injections, adult females aged 3 or 11 days after eclosure were injected with a suspension of fluorescent beads (FluoSpheres carboxylate-modified 0.2 μm; Life Technologies); 50-69 nl of a 1:10 dilution in PBS of was injected as described previously (Makhijani et al., 2011). Flies were incubated at 25ºC for 30min-1h, followed by imaging, hemocyte releases or embedding for cryosectioning. For quantification of bead-positive hemocytes, images were taken with a Leica DMI4000B microscope followed by manual counting.

For bioparticle injections, adult females aged 6 to 11 days after eclosure were injected with fluorescent bioparticles (*E. coli* K-12 Strain Bioparticles TexasRed Conjugate; Invitrogen) or pH sensitive fluorescent bioparticles (pHrodo *E. coli* Bioparticles Conjugate for Phagocytosis; Invitrogen). Each fly was injected with 27.6-32.2 nl of bioparticles (1mg/100µl) and then incubated at 25º C for 4 hours before analysis. For ex vivo hemocyte analysis of differential body sections, flies were split into head, thorax, abdomen using a scalpel, individually placed into wells of 20 µl of PBS, and hemocytes were released from each section by poking and crushing. Hemocytes were imaged as described under Microscopy.

### Microscopy

Standard fluorescence images were obtained on a Leica DMI4000 microscope. A Leica M205FA stereomicroscope with motorized stage and Leica LAS Montage module was used to image hemocytes labeled by fluorescent reporters in live CO2-anesthetized flies or whole mount stainings. Immunostained adult cryosections and dissected preparations were imaged using a Leica SP5 confocal microscope.

### Hemocyte quantification from adult flies

To determine hemocyte numbers by visualization of fluorescent reporter expressing hemocytes through the cuticle, the number of GFP labeled hemocytes in thorax, three legs and head of one side of the animal was counted from montage images (described above) for 1, 7, 19, 24, 37 and 63 days old adults. For each time point, 4-10 males and 4-10 females were used to determine the mean value of number of hemocytes and standard deviation. For hemocyte quantification in the dorsal thorax/abdominal area in (Fig. 5C-F’), see ‘Fucci analysis and quantification of hemocytes’. To determine total hemocyte numbers by release, single flies were CO2-anesthetized, and dissected in a glass well slide containing defined amount of 70µl of Schneider’s *Drosophila* medium (Millipore). Under a fluorescence stereomicroscope, the head was removed and the ventral side of the head was carefully torn between the eyes, and subsequently pinched with forceps until all of the hemocytes were released. The empty head was then removed from the well. Next, the abdomen was torn open and, starting from the posterior end, squeezed and poked until the majority of hemocytes were released. The thorax was squeezed/crushed multiple times to release the remaining hemocytes. The carcass was transferred out of the well. 10µl of this hemocyte suspension was pipetted into a hemocytometer, and cells were counted under a fluorescence microscope. Cell numbers from four 1mm^2^ squares were averaged, and the total number of hemocytes per fly was calculated. A total of 10 females plus 2 males per time point were quantified which were processed in groups of 4 animals. Average number of hemocytes/fly from 12 animals and standard deviation were calculated for each time point.

### Bacterial infection

Animals for hemocyte MARCM, PermaTwin MARCM or EdU based proliferation experiments and qPCR to detect AMP expression were immune challenged by either injection or feeding of bacterial strains as outlined below.

For injections, bacterial cultures were grown overnight in LB broth with suitable antibiotics. The following morning, a 1:20 culture dilution was incubated at 37°C for 3 hours, OD measured (NanoDrop 2000c spectrophotometer, Thermo Scientific) while typically <1, and the culture was spun down and the resulting pellet resuspended in PBS to achieve specific calculated ODs as summarized below. Bacterial strains (kind gifts from Bruno Lemaitre laboratory) and ODs were as follows; *Escherichia coli* OD 3 or 6 (see figure legends); *Micrococcus luteus* OD 1.5 or 2; *Erwinia carotovora carotovora 15* (Lemaitre et al., 1997) OD 1.5; *Enterobacter cloacae* (ß12, Jean Lambert(Lemaitre et al., 1997)) OD 2 or 4 (see figure legends). Bacterial suspension or control (sterile PBS) was injected into the thorax or abdomen of anesthetized females aged 5-7 days using a Nanoject II injector (Drummond Scientific Inc.) fitted with a pulled glass capillary, using volumes of 9.2nl to 27.6nl as indicated; experiments with *btl-GAL4* used 1-2 days old flies. Infected adults were incubated at 29°C, or as indicated. F1 crosses of transgenes regulated by the temperature-sensitive *tub-GAL80*^*ts*^ (McGuire et al., 2003) were initially maintained at 18ºC, and then induced by shifting animals to 29ºC, starting 24 hours before injection for crosses with *HmlΔ-GAL4, UAS-GFP; tub-GAL80*^*ts*^, or 48 hours before injection for crosses with *btl-GAL4, UAS-GFP, tub-GAL80*^*ts*^. Immune challenge by feeding of *S. marcescens* was performed as described previously (Nehme et al., 2007). *S. marcescens* grown overnight from a single colony inoculated into 5 ml LB broth with 50ug/ml ampicillin was diluted to OD600 of 0.1 in 10 ml of fly food. At least 10 adults per cohort were used.

### qRT-PCR

Adult flies were frozen at −80ºC after incubation at 29ºC following injections (see figure legends for specific incubation times). Frozen flies were homogenized in TRIzol (Invitrogen), and RNA was isolated using Direct-zol RNA MiniPrep kit, following the manufacturer’s instructions (Zymo). 1 µg of RNA was reverse transcribed using iScript cDNA synthesis kit (BioRad) and the resulting cDNA was diluted 1/10 for the qPCR. qPCR was run in a BioRad CFX96 Touch Real-Time PCR System, or ABI Viia7 Real-Time PCR System using the BioRad iTaq SYBR Green Supermix. Primers used for qPCR are summarized in Supplemental Table 1.

### Author Contributions

This study was made possible by the combined talent of all authors over the course of more than ten years. PSB expertly planned, performed and statistically analyzed the majority of qPCR experiments, added the aspect of Upd3/Jak/Stat signaling, contributed some survival data, and majorly revised qPCR figures. KM expertly planned and generated complex *Drosophila* genotypes and planned, performed and analyzed lineage tracing, MARCM, EdU, hemocyte quantification and marker experiments, and provided key information on the *Drosophila* respiratory epithelia. LH creatively developed new ways to visualize *Drosophila* adult respiratory epithelia, and planned, performed and analyzed adult fly dissections, confocal microscopy, microbead and bioparticle localization, phagocytosis assays, qPCR and survival experiments. KG planned, performed and analyzed experiments related to tissue colocalizations, microbead localization, confocal imaging, EdU incorporation, reporter and marker analyses, qPCR experiments, and contributed to the writing of the manuscript. RB planned, performed and analyzed whole and split-fly hemocyte counts and –qPCRs, and other qPCR and initial survival experiments. BA developed and performed fly cryosectioning, and performed EdU, lineage tracing, dye and marker experiments. KK performed and analyzed survival experiments. SC performed and analyzed experiments related to bioparticle injections and pHrodo counts. DO performed survival experiments. CW performed fly sectioning and staining. KW and FG generated and contributed valuable *Fucci* transgenic lines, and expert hemocyte quantification, Fucci analyses and images. CR and EM kindly contributed the PermaTwin system prior to its publication. EJVR and BL trained KB in *Drosophila* infection techniques and assays, and generously provided wide-reaching expert advice and materials. KB conceived the study, guided lab members, discovered the anatomical link of hemocytes and respiratory epithelia, planned experiments, analyzed data, generated figures, and wrote the manuscript. All authors provided input on the manuscript.

## Acknowledgements

We thank I. Ando, U. Banerjee, E. Bach, M. Crozatier, J.P. Dudzic, C. Evans, A. Giangrande, L. Kockel, C. Kocks, T. Kornberg, M. Meister, P. Rao, B. Stramer, S. Younger, the Bloomington Stock Center, the DGRC and the VDRC for fly stocks and antibodies. We especially thank Prashanth Rao and Chrysoula Pitsouli for their expert advice on the *Drosophila* tracheal system and respiratory epithelia, and Mark Krasnow for expert advice on insect literature. KSG was supported by an American Heart Association fellowship. KM was supported by a Human Frontier Science Program long-term fellowship. PSB was supported by a Swiss National Science Foundation early postdoc mobility fellowship. K. J Woodcock and F. Geissmann were supported by a Wellcome Trust Senior Investigator Award (WT101853/C/13/Z) to FG. This work was supported by grants from the American Cancer Society RSG DDC-122595, American Heart Association 13BGIA13730001, National Science Foundation 1326268, National Institutes of Health 1R01GM112083-01 and 1R56HL118726-01A1 (to KB). This investigation was in part conducted in a facility constructed with support from the Research Facilities Improvement Program, Grant number C06-RR16490 from the NCRR/NIH. Special thanks from KB to Steep Ravine Cabins at Mt. Tam State Park, CA.

## Supplemental Information

### Supplemental Experimental Procedures

#### *Drosophila* strains and fly husbandry

*Canton S, w1118*, or *yw* were used as control strains to match the background of experiment crosses. Transgenic lines and mutants used were *HmlΔ-GAL4* (Sinenko and Mathey-Prevot, 2004), *HmlΔ-DsRed* (Makhijani et al 2011), *HmlΔ-DsRednls* (Makhijani et al 2011), *srpHemo-GAL4* (Brückner et al. 2004), *srpGal4,UAS-lifeact-GFP* (from B. Stramer), *UAS-CD4-GFP; srpD-GAL4 (from M. Meister), btl-GAL4,UAS-GFP* (Rao et al., 2015) (from P. Rao), *ppl-GAL4* (Bloomington 58768), *Ubi-GAL4/Cyo* (Bloomington 32551). *UAS-GFP* (Song et al., 2007), *UAS-CD8GFP* (from L. Kockel), *UAS-srcEGFP* (from E. Spana), *UAS-Drosocin RNAi* (VDRC 42503, GD line on chr. 2, here labeled ‘D1’), *UAS-Drosocin-RNAi* (VDRC 105251, KK line line on chr. 2, here labeled ‘D2’), *UAS-imd RNAi/TM3, sb*^1^ (Bloomington 38933), *UAS-PGRP-LC* RNAi (Bloomington 33383), *UAS-upd3* RNAi (Bloomington 28575), *UAS-hop* RNAi (Bloomington 31319), *UAS-Stat92E* RNAi (Bloomington 31317) (all Bloomington TRiP RNAi lines correspond to y[1] v[1]; P{y[+t7.7]v[+t1.8]=TRiP.insertion}attP2), *UAS-imd* (Georgel et al., 2001) (from N. Buchon), *UAS-upd3* (from E. Bach), *UAS-hop*^*TumL*^ (Harrison et al., 1995) (from E. Bach), *UAS-3HA-Stat92E; UAS-3HA-Stat92E*^*ΔNΔC*^ (Ekas et al., 2010) (from E. Bach), *tub-GAL80*^*ts*^ (McGuire et al., 2003) (Bloomington 7018), *UAS-hid;UAS-rpr* (Zhou et al., 1995), *w^1118^;HmlΔFucciOrange*^*G1*^*;HmlΔFucciGreen*^*G2/S/M*^ (K. J Woodcock and F. Geissmann, see below); *Drosocin-GFP* (Tzou et al., 2000) (from B. Lemaitre), *Rel*^*E20*^ (Leulier et al., 2000) (from B. Lemaitre), *spz*^*rm7*^ (De Gregorio et al., 2002) (from B. Lemaitre). Lines for hemocyte MARCM *hsflp,UAS-GFP; HmlΔ-GAL4; FRT82B, tub-GAL80*, and *HmlΔ-GAL4, UAS-GFP; FRT82B* (Makhijani et al 2011) (experiment), and *HmlΔ-GAL4, UAS-GFP* (control). Lines for PermaTwin MARCM *w; FRT40A, UAS-CD8-GFP, UAS-CD2-Mir/Cyo; actGAL4, UAS-flp/TM6B* and *w; FRT40A, UAS-CD2-RFP, UAS-GFP-Mir/Cyo; tub-GAL80*^*ts*^ */TM6B* (Fernandez-Hernandez et al., 2013). Lines for lacZ lineage tracing: *tub-GAL80*^*ts*^*; UAS-Flp* and *srpHemo-GAL4, UAS-srcEGFP; Act>stop>nuc-lacZ* (Makhijani et al., 2011), or *HmlΔ-GAL4, UAS-GFP* and *UAS-Flp; tub-GAL80*^*ts*^ */CyO wee p; act>stop>lacZnls* (kind gift from P. Rao, Kornberg lab). Recombinant chromosomes and combinations of transgenes were generated by standard genetic techniques. Unless stated otherwise, all genetic crosses were kept at 25°C. Flies were raised on a standard dextrose cornmeal diet supplemented with dry yeast.

### HmlΔFucci transgenic lines

To generate *HmlΔFucci-G1-Orange* and *HmlΔFucciGreenS/G2/M-Green* constructs, Fucci cloning vectors (MBL laboratories, AM-V9014 and AM-V9001) were both digested with HindIII and BamHI in order to excise the insert Fucci fluorescence genes; *Fucci-G1 Orange* and *Fucci-S/G2/M Green*. A previously cloned *p-Red H-Stinger-HmlΔ-DsRed* plasmid (Clark et al., 2011) was digested with BamHI and SpeI in order to excise *DsRed* whilst leaving the *HmlΔ* promoter in place. T4 DNA polymerase was used o blunt the *p-Red H-Stinger-HmlΔ* vector and the *Fucci* DNA inserts. Vector and *Fucci* insert DNA were then digested again with BamHI for a sticky-blunt ligation (Takara Bio Inc -TaKaRa DNA Ligation Kit LONG). Cloning was done according to standard procedures. The separate *HmlΔFucci-G1-Orange* and *HmlΔFucciGreenS/G2/M-Green* constructs were injected into *w^1118^ Drosophila* embryos to generate transgenic lines (BestGene Inc.). Plasmid maps and more detailed cloning information are available upon request.

### Flipout-LacZ lineage tracing

Flipout-lacZ lineage tracing was essentially done as described by (Weigmann and Cohen, 1999). Lineage tracing of embryonic hemocytes was described previously in (Makhijani et al., 2011). F1 progeny of the following cross were used *tub-GAL80*^*ts*^*; UAS-Flp x srphemo-GAL4, UAS-srcEGFP; Act>stop>nuc-lacZ*. To obtain specific lacZ labeling of embryonic hemocytes, expression of *srpHemo-GAL4* through *tub-GAL80*^*ts*^ was controlled by shifting 5 hour egg collections from 18°C to the permissive temperature of 31°C for 6 hours and then back to 18° C until pupal or adult stage. The no heat shock control was continuously maintained at of 18°C. For lineage tracing of embryonic-lineage hemocytes from the larva, F1 progeny of the following cross were used *HmlΔ-GAL4, UAS-GFP x UAS-Flp; tub-GAL80*^*ts*^ */ CyO wee p; act>stop>lacZnls*. Heat shock at 29°C was administered from 0-48h AEL, and animals were then maintained at 18°C for the remainder of their life. This time window was chosen to avoid labeling of lymph gland-lineage hemocytes that express the *HmlΔ-GAL4* driver from a later point during larval development. βGal positive hemocytes, relative to all Crq positive plasmatocytes, were determined by immunostaining of ex vivo released hemocytes at two points, (1) from 3^rd^ instar larva before mobilization of differentiated lymph gland hemoytes takes place, and (2) from adult animals at 12 days post eclosure. The relative fraction of labeled hemocytes in the adult compared to the larva was calculated from the average and standard deviation from 16 adult females.

### Fucci analysis and quantification of hemocytes

Fucci analysis, imaging and external hemocyte counts for Fig. 3D and 5C-F’ were performed using a Leica SP5 confocal microscope. Flies were affixed dorsally to glass cover slips with superglue (Loctite) Sf435746 and imaged in vivo using 20x Dry NA 0.5 or 40x Oil NA 1.25 objectives. Cohorts of 15-21 flies per condition were imaged, and images were processed using Fiji and Imaris software. Cell counts were performed using the MATLAB spot detection function in Imaris, calculating the mean±SEM, and performing an unpaired t test to calculate statistical significance. For bacterial infections related to Fucci analyses and hemocyte counts, individual *E. coli, M. luteus*, or *E. cloacae* cultures were grown at 37°C overnight. The following morning, cultures were centrifuged at 4°C at 1600rpm for 10 minutes. Bacterial pellets were re-suspended in PBS to a final OD600 of 1. Injection was performed using a PicospritzerR III (Parker Hannifin), and the injection volume was calibrated to 50nl by injecting a drop into a plot of oil. Adult males 7 days post eclosion were used for injections of bacteria, sterile PBS, and non-injected controls.

### Hemocyte MARCM

We developed Hemocyte MARCM as a derivation of the previously described MARCM (Lee and Luo, 1999), a labeling method based on mitotic recombination. The ubiquitous *tub-GAL4* driver was replaced with *HmlΔ-GAL4* to allow specific GFP labeling of all dividing cells that would ultimately show Hml+ plasmatocyte characteristics (Makhijani et al., 2011). Genotypes of F1 progeny were as follows: Hemocyte MARCM: *hsflp,UAS-CD8GFP/ +; HmlΔ-GAL4/ HmlΔ-GAL4, UAS-GFP; FRT82B, tub-GAL80/ FRT82B*. Control: *hsflp, UAS-CD8GFP/ +; HmlΔ-GAL4/ HmlΔ-GAL4, UAS-GFP; FRT82B, tub-GAL80/ +*. Various heat shock and incubation schemes were tested to optimize Hemocyte MARCM labeling, using 1-2 week old adult progeny. Heatshock was induced at 37°C for 1h 3 times/day for 3 days; 30°C continuously for 4 days; 32°C continuously for 4 days. Following heat shock schemes, flies were maintained at room temperature and observed under a fluorescence stereomicroscope for GFP labeled hemocytes. The control cross lacks one of the FRT82B chromosome and therefore does not allow Flp-induced recombination to occur. Progeny from control crosses was also exposed to heat shock, allowing to observe non-specific GFP labeling due to heat shock conditions and imbalance of GAL4 and GAL80, and/or occasional random recombination events. Groups of 12-18 animals (equal mix of males and females) were examined per condition. For immune challenges, 4-6 days old flies were infected with a mixture of *E. coli* and *M. luteus*, by pricking flies in the thorax with a needle dipped in a bacterial pellet (mix of equal OD600 from overnight cultures of *E. coli* and *M. luteus*).

### PermaTwin MARCM

For PermaTwin MARCM, genotypes were essentially as described in (Fernandez-Hernandez et al., 2013). F1 progeny of the following cross were used: *w; FRT40A, UAS-CD8-GFP, UAS-CD2-Mir/Cyo; actGAL4, UAS-flp/TM6B* x *w; FRT40A, UAS-CD2-RFP, UAS-GFP-Mir/Cyo; tub-GAL80*^*ts*^*/TM6B*. Flies were raised at 18°C until adulthood to repress *flp* expression. Adult progeny of different ages (3-5 days and 2-3 weeks, n=12/age and induction condition) were examined. Animals were shifted to 29°C for varying time windows of 4 days, 8 days, 2 weeks, 3 weeks, respectively, and observed by live imaging for appearance of GFP and RFP labeled hemocytes; control tissues with labeled clones were dissected and observed in parallel. Adults were also dissected in PBS, to examine for hemocyte populations that might be attached to internal tissues. Negative control flies were maintained at 18°C continuously. In order to test the effect of immune challenge on hemocyte proliferation in adults, flies of the above age and genotyopes were infected with either *E. coli* (OD 3) or *M.luteus* (OD 1.5 or 2) and observed 2-8 days post-infection.

### Quantification of AMP expression by qPCR

For qPCR analysis, total RNA was harvested from 10 pooled adult female *Drosophila* for each experimental condition. For qPCR on split fly samples, 8 pooled females for each condition were snap frozen in LN2, and for each fly the head/thorax and abdomen were separated using a scalpel on a metal block on dry ice. Frozen body parts or whole flies were pestle-homogenized in 500 μl Trizol reagent (Millipore), and RNA was purified using a Direct-zol RNA Miniprep kit (Zymo Research). RNA was quantified and quality checked with a NanoDrop 2000c spectrophotometer (Thermo Scientific). 1 μl of total RNA was used in a 20 μl iScript cDNA Synthesis (BioRad) reverse transcription reaction. qPCR was carried out on a CFX96 c1000 Thermal Cycler (BioRad) using IQ SYBR Green Super Mix (BioRad) for *Drosocin, Drosomycin, Diptericin, Cecropin A1*, and *Attacin A*. Relative mRNA expression levels were normalized to that of *RpL32*; for primers see Supplemental Table 1. Results are presented as the mean and standard deviation of at least three (in rare cases two) biological repeats.

**Supplemental Table 1.**
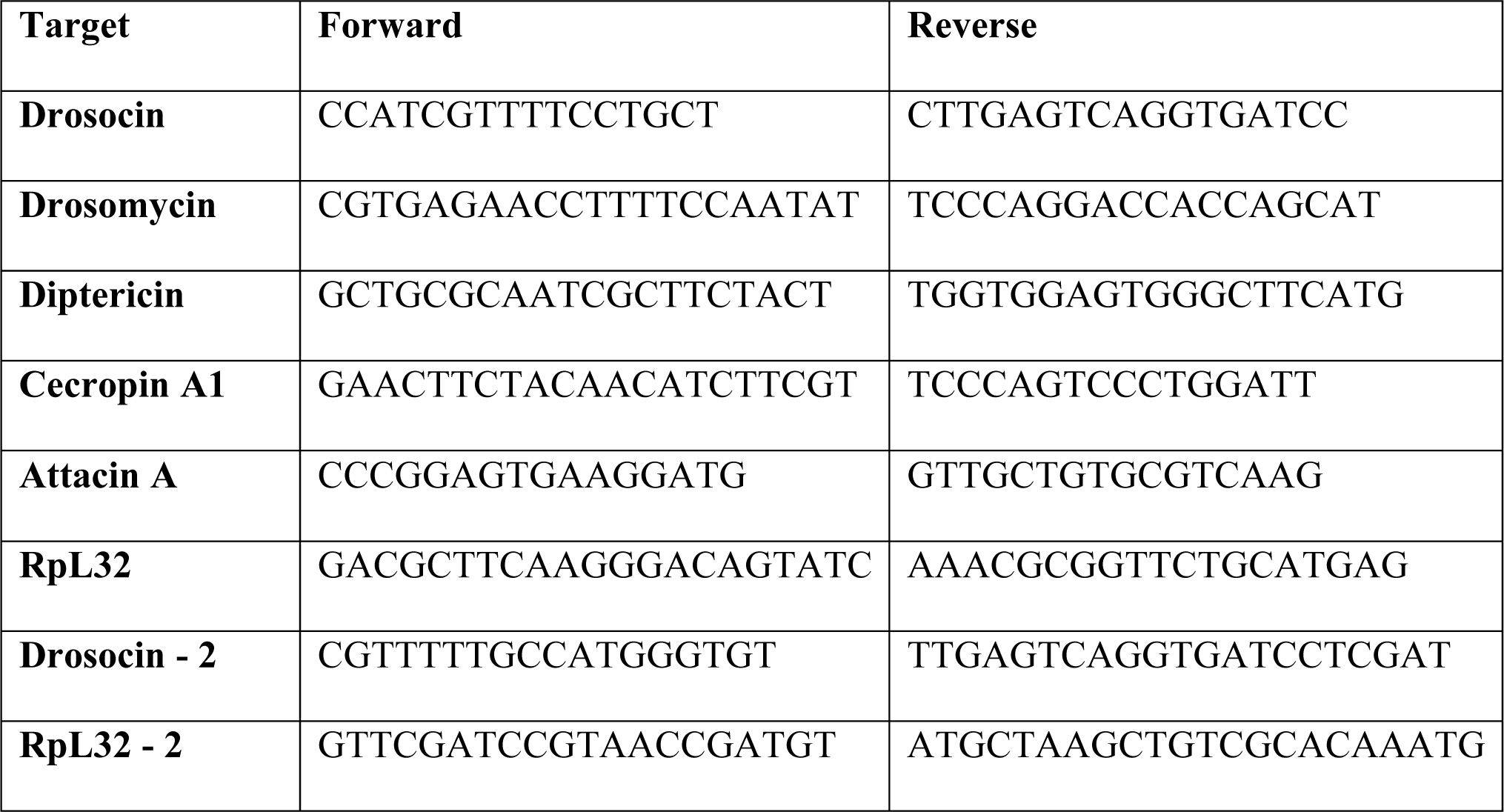
Primer sequences.

## Supplemental Figures

**Supplemental Figure 1.**
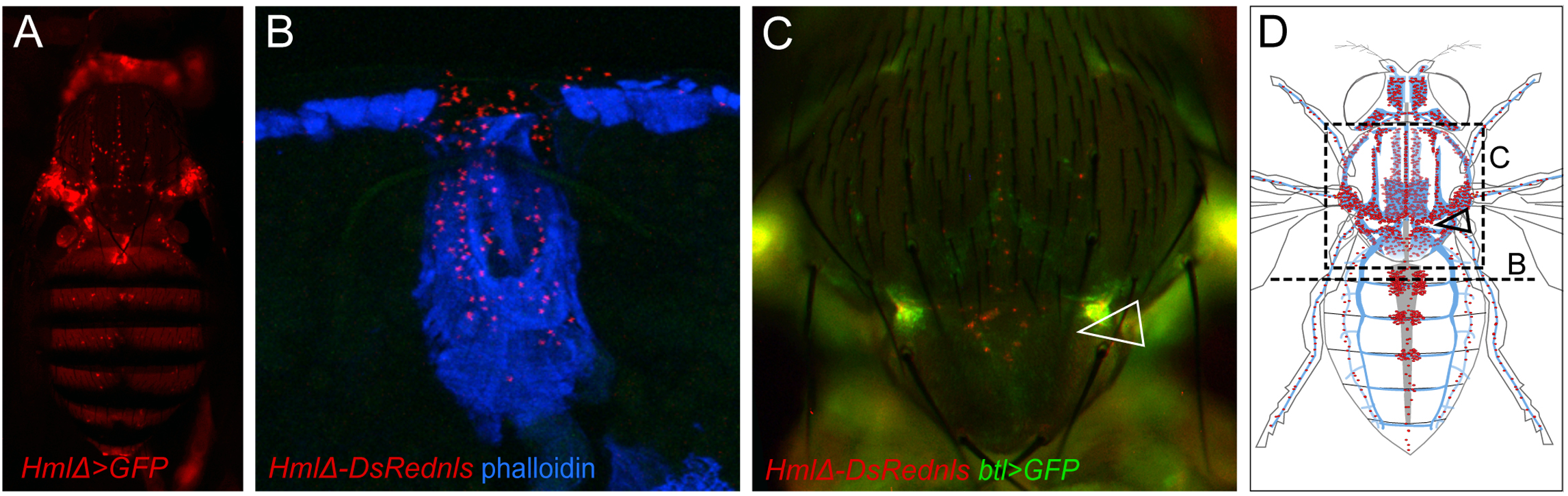
The respiratory reservoir of hemocytes, and hemocyte clusters at the ostia of the heart. (A) *Drosophila* adult, live imaging *HmlΔ-GAL4, UAS-GFP* (hemocytes red pseudocolor to match model), anterior up. (B) Cross section through the anterior abdomen, *HmlΔ-DsRednls* (hemocytes, red), phalloidin blue, dorsal up. (C) Thorax of adult *Drosophila*, live imaging of intact adult *HmlΔ-DsRednls*/*btl-GAL4, UAS-GFP* (hemocytes red, tracheal system green); arrowhead points to scutellum where hemocytes accumulate along the respiratory epithelia. (D) Schematics of heart (grey), respiratory epithelia of the thorax and head and other parts of the tracheal system (blue); associated hemocytes (red). Dashed lines indicate section shown in B, and area shown in C, arrowhead points to corresponding area in (C).

**Supplemental Figure 2.**
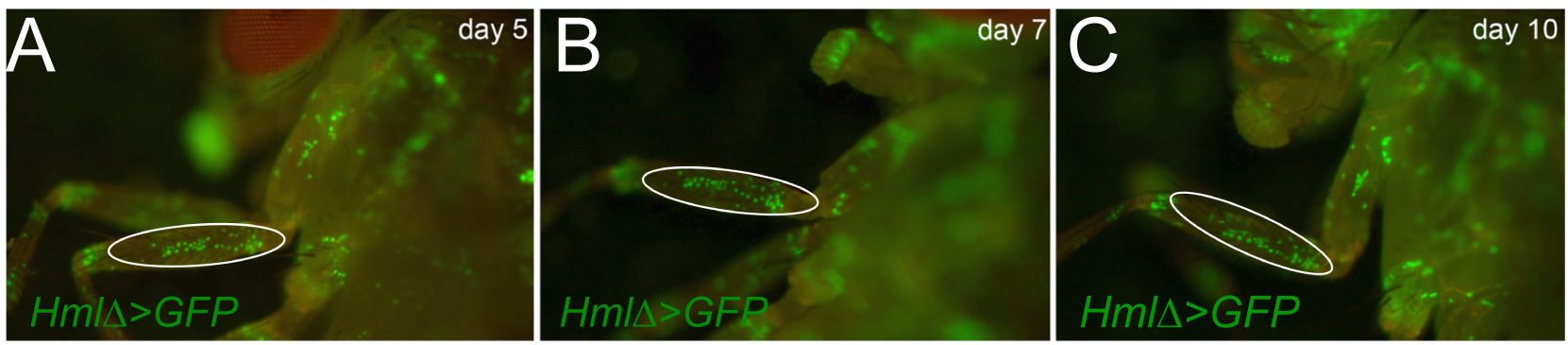
Hemocytes are stationary; increased expression of *Hml* and *Crq* post infection. (A-C) Time course of resident hemocyte pattern, live imaging of representative animal, genotype is *HmlΔ-GAL4, UAS-GFP*; day 5, 7, 10 post eclosure.

**Supplemental Figure 3.**
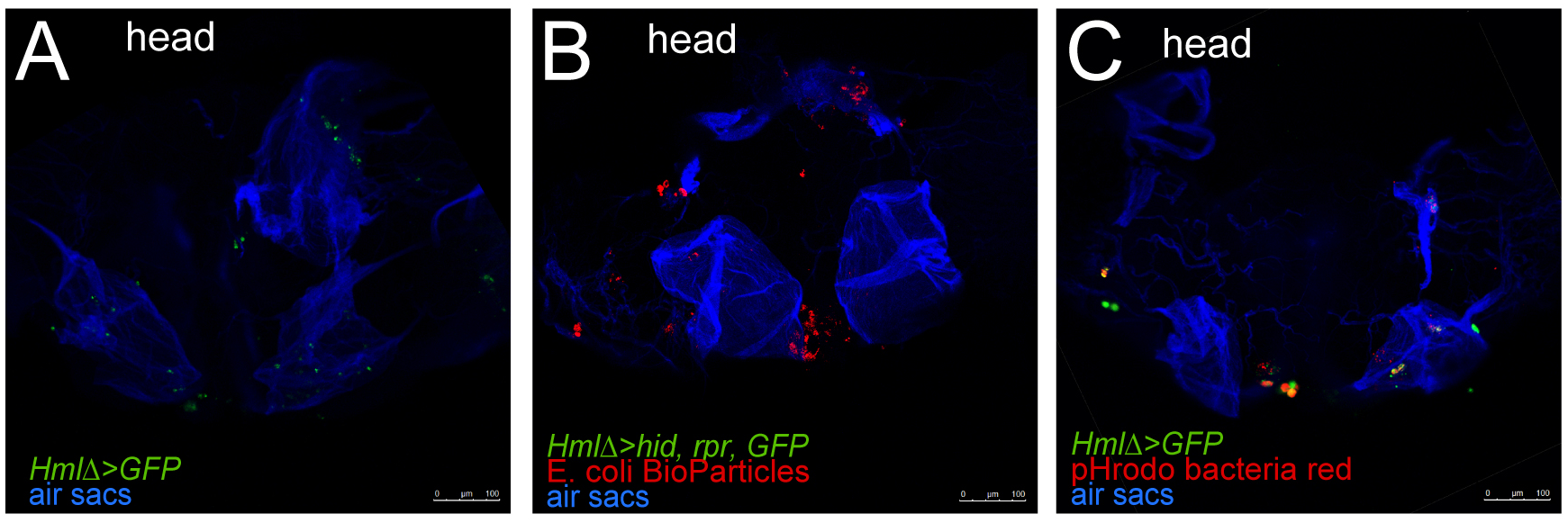
Bioparticle accumulation at the respiratory epithelia. (A-C) Confocal imaging of respiratory epithelia, hemocytes, and fluorescent bioparticles in dissected head preparations. (A) Control genotype *Hml-GAL4, UAS-GFP/+*, no bioparticle injection; hemocytes green, respiratory epithelia blue. (B) Hemocyte ablation genotype *Hml-GAL4, UAS-GFP/UAS-rpr; UAS-hid/+, E. coli* (K-12 strain) bioparticles, TexasRed, injected (1mg/100ml PBS; 32.2nl); hemocytes green (absent), bioparticles red, respiratory epithelia blue. (C) Control genotype *Hml-GAL4, UAS-GFP/+*, pHrodo red *E. coli* bioparticles injected (1mg/100µL PBS; 32.2nl); hemocytes green, bioparticles red, respiratory epithelia blue.

**Supplemental Figure 4.**
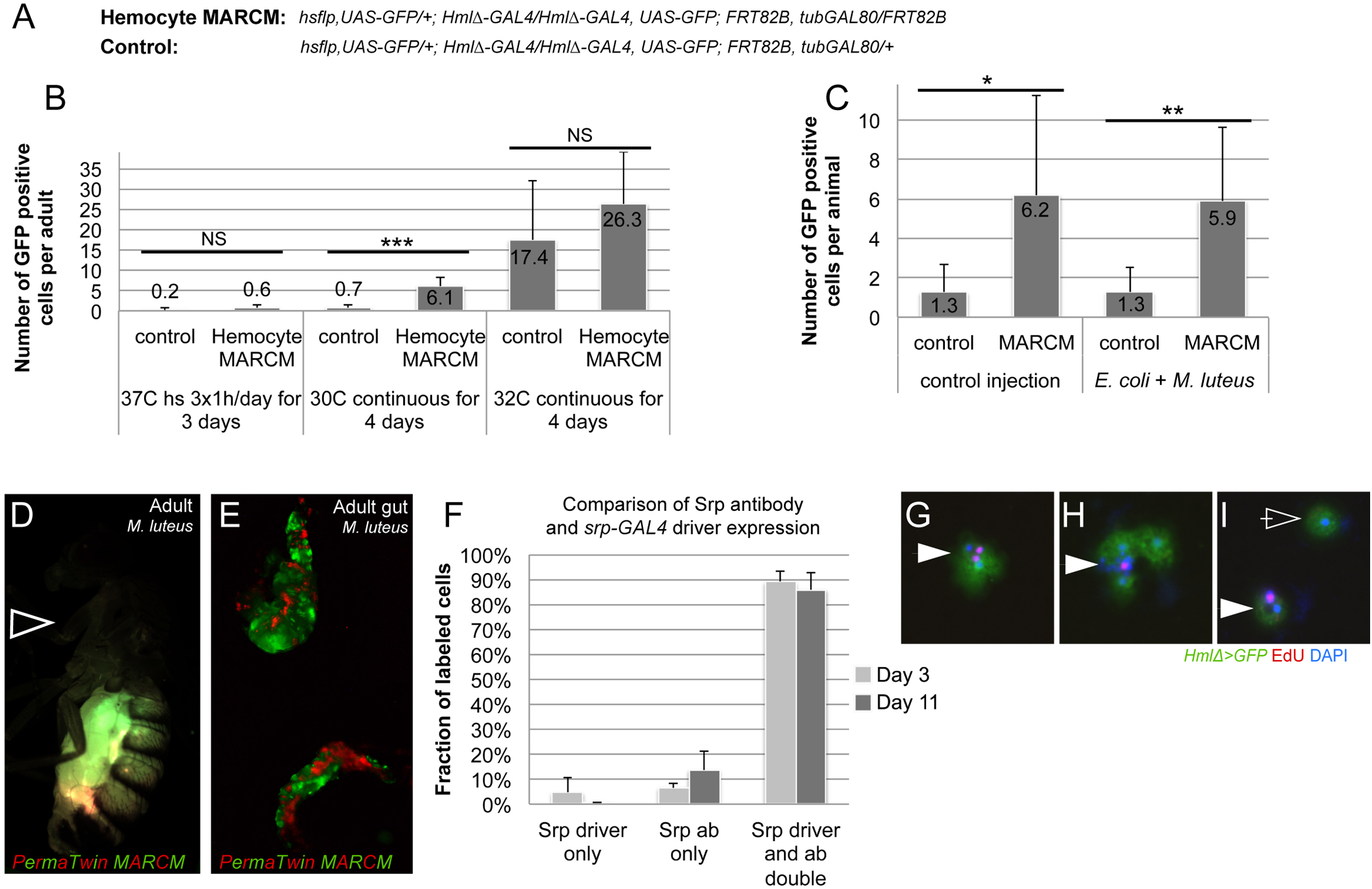
No significant new production of hemocytes in adult *Drosophila*. (A-C) MARCM clonal labeling to detect any dividing cells that eventually give rise to hemocytes. (A) genotypes of F1 progeny used; (B) titration of conditions for MARCM induction; (C) at 30°C 4 days continuous induction, experiment animals show at best an average of 6 labeled hemocytes over background; this number is not enhanced upon bacterial infection. (D, E) PermaTwin 2-color clonal analysis to search for dividing cells that would give rise to hemocytes; genotype of F1 progeny used is *w^1118^; FRT40A, UAS-CD8-GFP, UAS-CD2-Mir/ FRT40A, UAS-CD2-RFP, UAS-GFP-Mir; actGAL4, UAS-flp/ tub-GAL80*^*ts*^. (D) Adult fly after 5 days of heat shock induction and infection with *M. luteus*; arrowhead points to thorax and head, which show no sign of hemocyte labeling. Representative images were taken from 3 independent biological repeats (n=8/repeat) of various time points and infection conditions. (E) positive control, dissected gut showing PermaTwin clones. (F) Comparison of labeling efficiency of *srpGal4, UAS-lifeact-GFP* and anti-Srp antibody staining, day 3 and 11 animals were examined. (G-I) Examples of hemocytes with EdU positive inclusions. Careful examination and counterstaining with DAPI revealed that all EdU labeled elements correspond to phagocytic vesicles (arrowheads); hemocyte nuclei always remained EdU negative (open arrowhead).

**Supplemental Figure 5.**
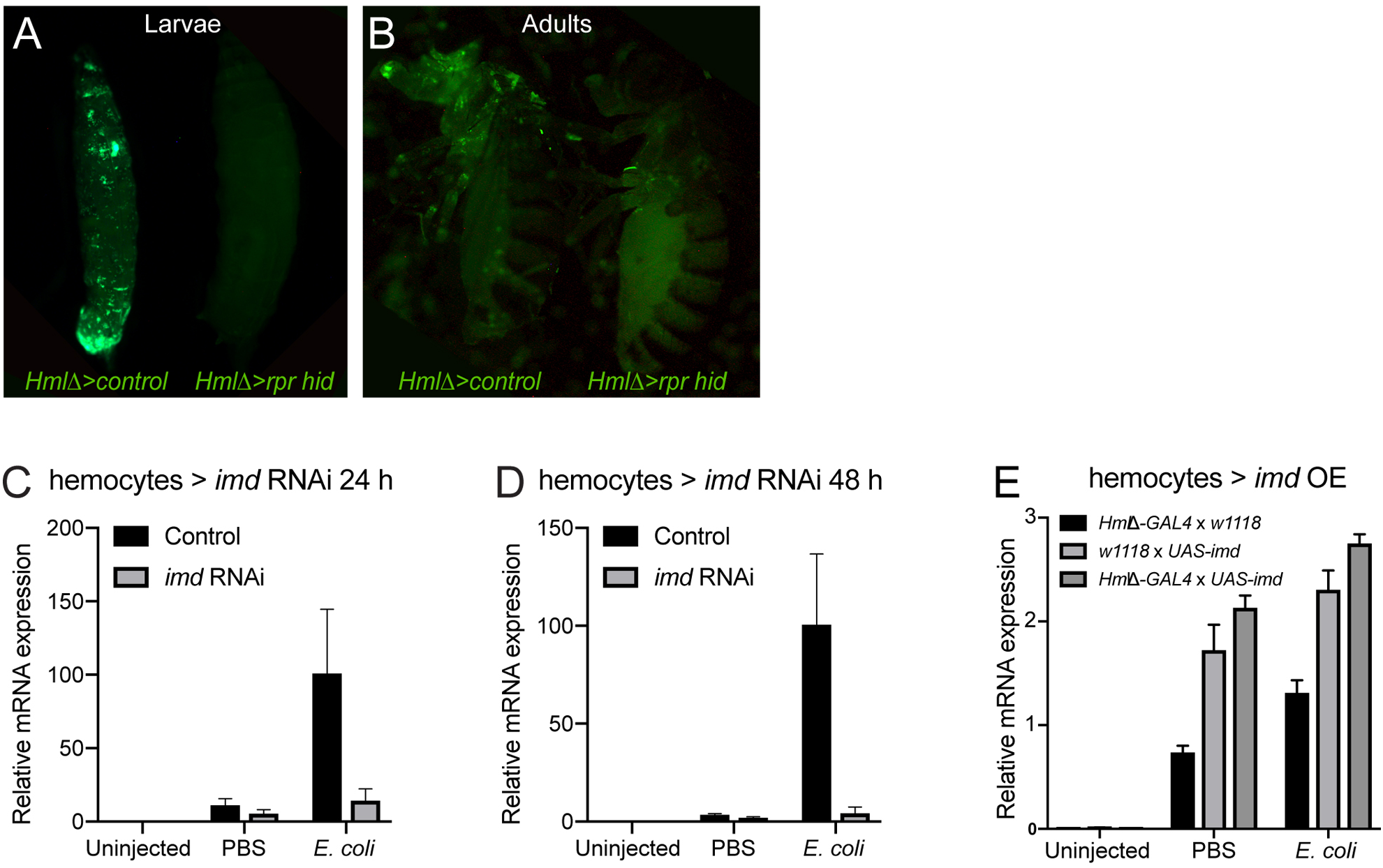
AMP gene expression depends on hemocytes; Imd signaling in hemocytes. (A, B) Live images of hemocyte-ablated animals and controls, hemocytes in green; genotypes are *HmlΔ-GAL4,UAS-GFP/+* (control, animal on left side) and *w; HmlΔ-GAL4,UAS-GFP/ UAS-rpr; UAS-hid/ +* (hemocyte ablation, animal on right side). (A) Larvae; (B) adults, lateral view. (C-D) *Drosocin* qPCR of whole flies, *imd* RNAi silencing in hemocytes. Genotypes are *HmlΔ-GAL4, UAS-GFP/+; UAS-imd RNAi*/+ (experiment) and *HmlΔ-GAL4, UAS-GFP/+* (control). 5-6 day old adult females were left untreated (control) or injected with PBS or *E.coli* in PBS (OD 3), 27.6nl. Flies were analyzed at 24h (C) and 48h (D) post infection. Each chart displays mean and SEM of samples from a representative biological replicate experiment, using pools of 10 females per condition, and triplicate qPCR runs. (E) *Drosocin* qPCR of whole flies, *Imd* overexpression in hemocytes, genotypes are control *HmlΔ-GAL4, UAS-GFP/+* versus *HmlΔ-GAL4, UAS-GFP/ UAS-imd*; 5 day old adult females were left untreated (control) or injected with PBS or of *E.coli* in PBS (OD 3), 27.6nl. Each chart displays mean and SEM of samples from a representative biological replicate experiment, using pools of 10 females per condition, and triplicate qPCR runs.

**Supplemental Figure 6.**
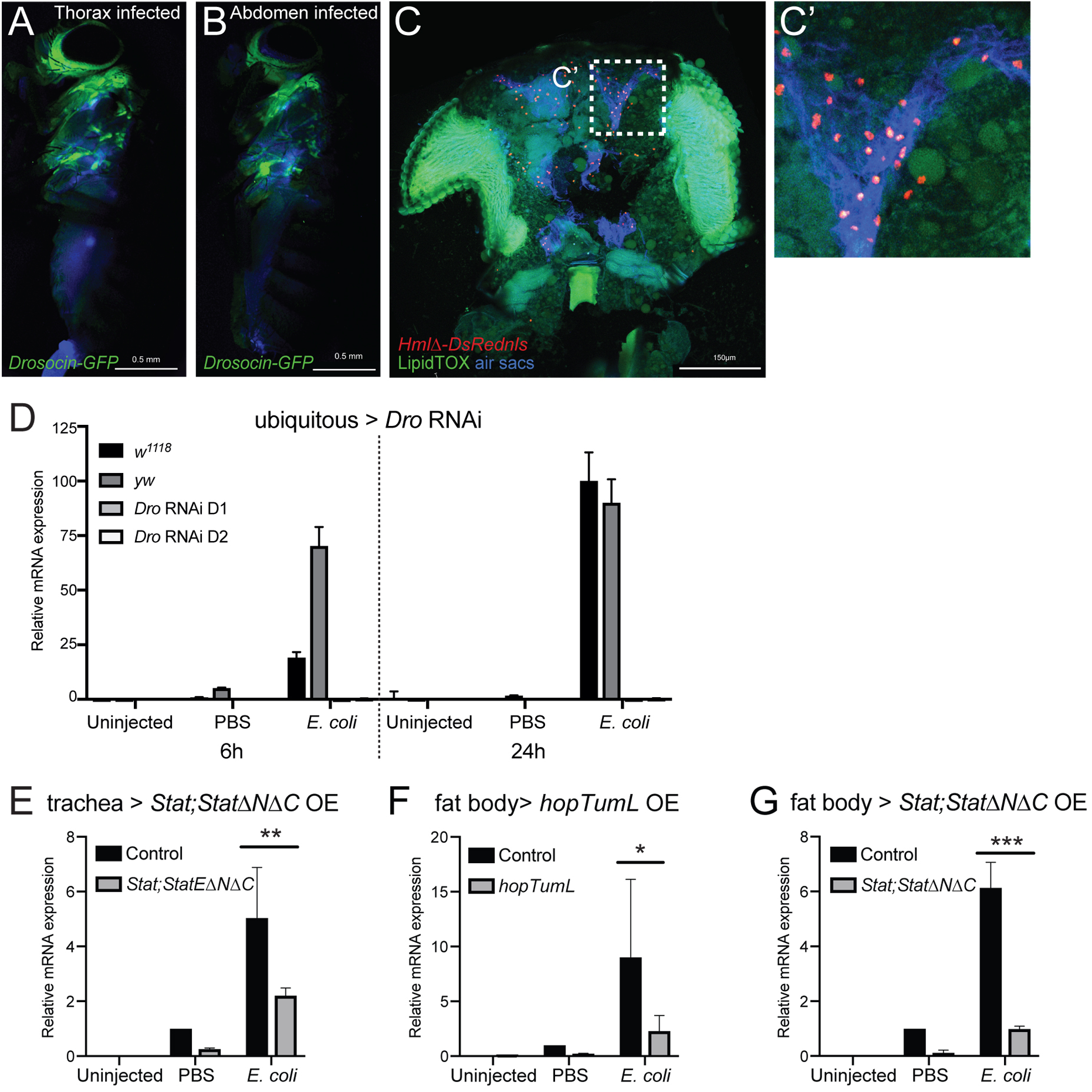
*Drosocin* expression in respiratory epithelia and fat body, and the role of Jak/Stat signaling. (A-B) *Drosocin-GFP* expression is found in head and thorax, independent of the site of infection. (A) *Drosocin-GFP* 2 days post thorax injection (*E. coli*), respiratory epithelia (air sacs, blue); (B) *Drosocin-GFP* 2 days post abdomen injection (*E. coli*), respiratory epithelia (air sacs, blue). (C-C’) Anatomy of fat body tissue lining the respiratory epithelia and hemocytes; *HmlΔ-DsRednls* (hemocytes, red), fat body (LipidTOX, green), respiratory epithelia (air sacs, blue) (C) Head cross section. (C’) Closeup of region indicated in (C). Note that hemocytes are layered between respiratory epithelia and fat body. (D) *Drosocin* qPCR of whole flies, ubiquitous RNAi knockdown of *Drosocin*; genotypes are *Ubi-GAL4/UAS-Drosocin* RNAi line D1 (GD) or *Ubi-GAL4/UAS-Drosocin* RNAi line D2 (KK) or control *Ubi-GAL4/*+ (cross with *yw* or *w^1118^*). 5 day-old females were left untreated or injected with PBS or *E. coli* in PBS (OD 6), 9.2nl, and harvested at 6, 12, 24 h post infection. Chart displays mean and SEM of samples from a representative biological replicate experiment, using pools of 10 females per condition, and triplicate qPCR runs. *Drosocin* kd efficiency is at 6h=98.2%, 12h=98.8%, 24h=98.9%. (E-G) qPCR of *Drosocin* in whole adult flies upon overexpression (OE) of Jak/Stat signaling components in the tracheal system or fat body. *Drosophila* untreated, injected with PBS, or with of *E.coli* in PBS (OD 6), 9.2 nl; flies were harvested at 6h post injection. Each chart displays the mean and CI of samples from 3 averaged biological replicate experiments, using pools of 10 females per condition, and triplicate qPCR runs for each sample. Values of all charts are displayed relative to the average RNA level induced by sterile PBS injections in control flies. Two-way ANOVA with Sidak’s multiple comparison test was performed, *,**,***,or **** corresponding to p≤0.05, 0.01, 0.001, or 0.0001, respectively (Prism). (E) *Drosocin* qPCR, genotype is control *btl-GAL4, tub-GAL80*^*ts*^, *UAS-GFP*/+ versus *UAS-3HA-Stat92E /+; UAS-3HA-Stat92E*^*ΔNΔC*^/ *btl-GAL4, tub-GAL80*^*ts*^, *UAS-GFP* (G-H) *Drosocin* qPCR, (G) genotype is control (*ppl-GAL4, UAS-GFP/+*) versus *ppl-GAL4, UAS-GFP*/ *UAS-hop*^*TumL*^; (H) genotype is control versus *UAS-3HA-Stat92E/ ppl-GAL4, UAS-GFP*; *UAS-3HA-Stat92E*^*ΔNΔC*^./+.

**Supplemental Figure 7.**
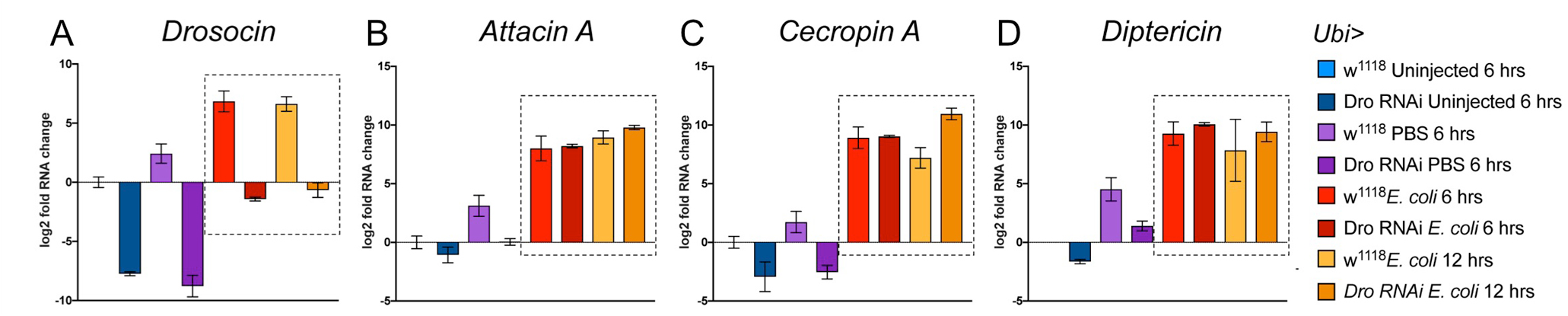
Endogenous *Drosocin* promotes survival after infection; *Drosocin* is not required for induction of other AMPs after infection. (A-D) Expression of AMP genes in the background of ubiquitous *Drosocin* silencing. Genotypes are *Ubi-Gal4*/*UAS-Drosocin* RNAi D1 *and Ubi-Gal4*/*UAS-Drosocin* RNAi D2, and control Ubi-GAL4/+ cross with *w^1118^*. Conditions are uninjured controls, injection of PBS and injection of *E. coli* in PBS (OD 6), 9.2 nl, with time points 6h and 12h post injection as indicated. Each chart displays the log2 mean and SEM of samples derived from pools of 10 females per condition, and triplicate qPCR runs. AMP genes quantified by qPCR for expression are (A) *Drosocin*; (B) *Attacin A*; (C) *Cecropin A1*; (D) *Diptericin*. The log2 mean is displayed to illustrate subtle expression changes below baseline. Note that *Drosocin* knockdown does not affect the induction of other AMPs following gram-negative infection, although it may mildly affect expression levels of other AMPs in uninjected condition, and under sterile injury (PBS injection).

